# Palmitoylation targets the Calcineurin phosphatase to the Phosphatidylinositol 4-kinase complex at the plasma membrane

**DOI:** 10.1101/2021.07.14.451536

**Authors:** Idil Ulengin-Talkish, Matthew AH Parson, Meredith L Jenkins, Jagoree Roy, Alexis ZL Shih, Nicole St-Denis, Gergo Gulyas, Tamas Balla, Anne-Claude Gingras, Péter Várnai, Elizabeth Conibear, John E Burke, Martha S. Cyert

**Author notes:** Max-Delbrueck Center for Molecular Medicine, Berlin, Germany (Alexis ZL Shih).

## Abstract

Calcineurin, the conserved protein phosphatase and target of immunosuppressants, is a critical mediator of Ca^2+^ signaling. To discover novel calcineurin-regulated processes we examined an understudied isoform, CNAβ1. We show that unlike canonical cytosolic calcineurin, CNAβ1 localizes to the plasma membrane and Golgi due to palmitoylation of its divergent C-terminal tail, which is reversed by the ABHD17A depalmitoylase. Palmitoylation targets CNAβ1 to a distinct set of membrane-associated interactors including the phosphatidylinositol 4-kinase (PI4KA) complex containing EFR3B, PI4KA, TTC7B and FAM126A. Hydrogen-deuterium exchange reveals multiple calcineurin-PI4KA complex contacts, including a calcineurin-binding peptide motif in the disordered tail of FAM126A, which we establish as a calcineurin substrate. Calcineurin inhibitors decrease PI4P production during Gq-coupled GPCR signaling, suggesting that calcineurin dephosphorylates and promotes PI4KA complex activity. In sum, this work discovers a new calcineurin-regulated signaling pathway highlighting the PI4KA complex as a regulatory target and revealing that dynamic palmitoylation confers unique localization, substrate specificity and regulation to CNAβ1.

## Introduction

Cells respond to changes in their environment via signaling pathways, including those regulated by calcium ions (Ca^2+^). The amplitude and duration of dynamic changes in the intracellular Ca^2+^ concentration provide specific temporal and spatial cues that direct a myriad of physiological responses. Hence, elucidating mechanisms that initiate Ca^2+^ signaling and identifying downstream Ca^2+^ sensing-effectors are critical for understanding cellular regulation in both healthy and diseased cells.

Calcineurin (CN/PP2B/PPP3), the conserved Ca^2+^/calmodulin (CaM)-activated serine/threonine protein phosphatase, transduces Ca^2+^ signals to regulate a wide-array of physiological processes. In humans, CN is ubiquitously expressed and has well-established roles in the cardiovascular, nervous, and immune systems ^1–3^. Because CN dephosphorylates NFAT (Nuclear Factor of Activated T-cells) transcription factors to activate the adaptive immune response ^4^, CN inhibitors FK506 (Tacrolimus) and cyclosporin A (CsA) are in wide clinical use as immunosuppressants ^5^. However, by inhibiting CN in non-immune tissues, these drugs also provoke a variety of unwanted effects which underscores the need to comprehensively map CN signaling throughout the body. Recently, systematic discovery of CN targets revealed that many CN-regulated pathways are yet to be elucidated ^6, 7^. Here, we uncover novel aspects of CN signaling by focusing on an understudied isoform, CNAβ1.

Calcineurin is an obligate heterodimer of catalytic (CNA) and regulatory (CNB) subunits. In mammals, three isoforms of CNA (α, β and γ) are encoded by separate genes with tissue specific expression. These isoforms display a similar domain architecture consisting of a catalytic domain, binding sites for CNB and CaM, and a C-terminal autoinhibitory domain (AID) which blocks phosphatase activity under basal conditions. Under signaling conditions that give rise to cytosolic Ca^2+^ levels, binding of both Ca^2+^ and Ca^2+^/CaM to CNB and CNA, respectively, disrupts inhibition of the catalytic site by the AID ^8–12^. This activation mechanism is conserved across all CN isoforms in animals and fungi, with the only known exception being a transcript variant of the CNAβ gene, termed CNAβ1 ^13–15^.

Alternative 3’ end processing of the *PPP3CB* mRNA gives rise to two CNAβ isoforms, CNAβ2 with canonical architecture, and the non-canonical CNAβ1 ^13, 14^. CNAβ1 and CNAβ2 share N-terminal sequence identity through the CaM-binding domain, but exclusion of two terminal exons and subsequent translation of intronic sequences results in a divergent C-terminus for CNAβ1 that is hydrophobic and lacks the AID (Fig. 1a). This alternative C-terminal tail is conserved in vertebrates (Fig. 1b) and CNAβ1 is broadly expressed in human tissues at a low level, alongside the canonical CN isoforms ^14, 15^. *In vitro* biochemical characterization of CNAβ1 identified an autoinhibitory sequence, ^462^LAVP^465^, in its C-terminal tail, which impedes substrate binding ^15^. CN recognizes substrates by binding two short, degenerate peptide motifs, “PxIxIT” and “LxVP”, found primarily in the disordered regions of its substrates ^16^. LxVP motifs bind to a region at the CNA/CNB interface that is accessible only after Ca^2+^/CaM binding ^16–21^. FK506 and CsA inhibit CN by blocking this LxVP binding pocket, showing that this interaction is essential for dephosphorylation ^21^. Notably, the maximal activity of CNAβ1 is limited compared to CNAβ2 due to this LxVP-mediated autoinhibition which is only partially relieved by Ca^2+^/CaM *in vitro* ^15^. However, mechanisms that govern the activity of this isozyme *in vivo* remain to be investigated.

**Figure 1:**
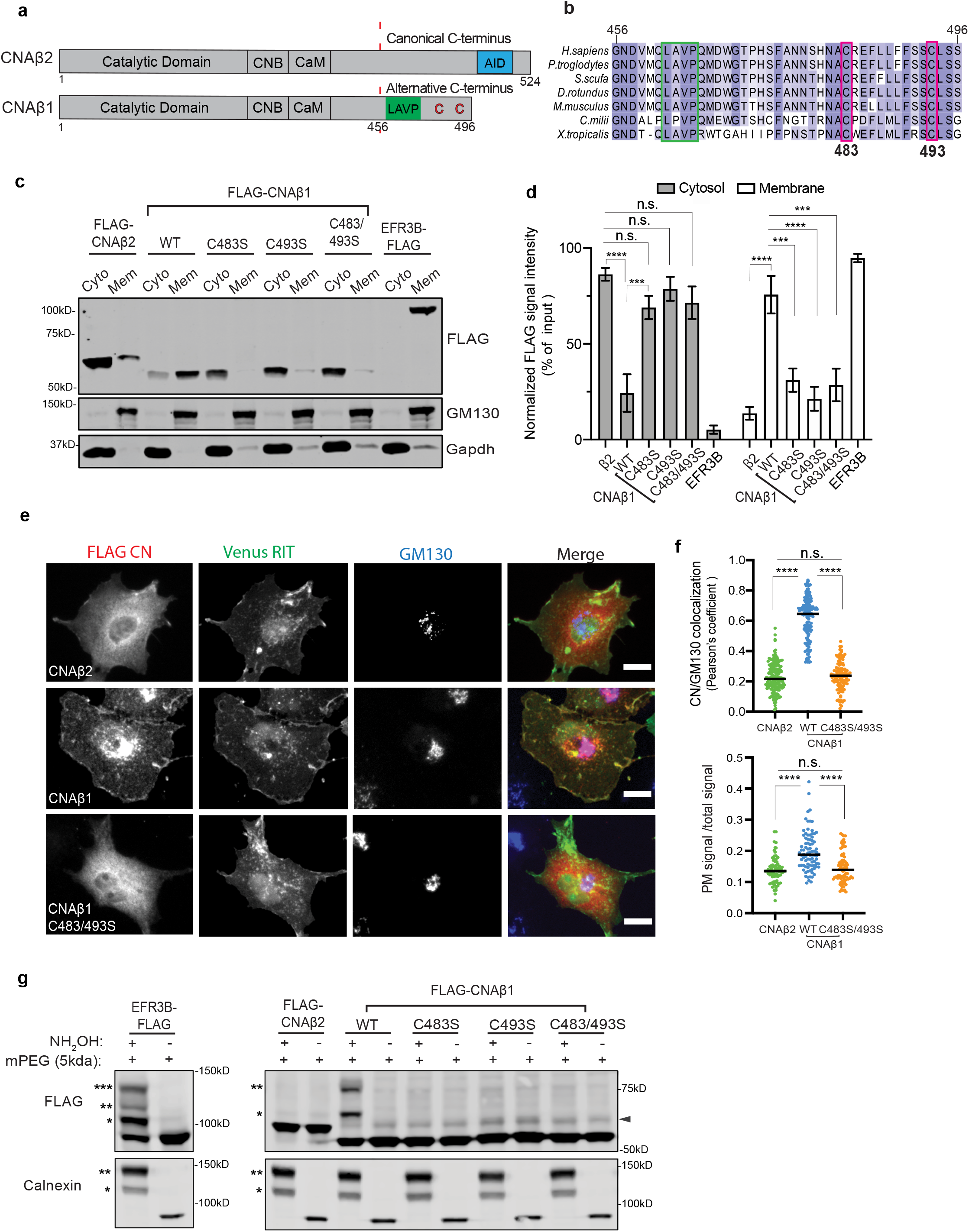
CNAβ1 localizes to intracellular membranes via palmitoylation at two conserved cysteines unique to its C-terminal tail. **a** Schematic of the domain architecture of CNAβ isoforms. Regulatory subunit binding domain (CNB), calmodulin binding domain (CaM). Autoinhibitory domain, AID, shown in blue; LAVP sequence shown in green. Palmitoylated cysteines are in red. **b** Sequence conservation of the CNAβ1 C-terminal tail (a.a 456-496) across vertebrates. The autoinhibitory LAVP sequence motif (green) and palmitoylated cysteines (red, C483 and C493), are boxed. **c** Representative immunoblot of subcellular fractions of COS-7 cells transfected with FLAG-CNAβ2, -CNAβ1 (WT or cysteine mutants) or EFR3B-FLAG. The CNAβ isoforms and EFR3B were detected using anti-FLAG antibody. GM130 and GAPDH were used as membrane and cytosol markers, respectively. **d** Quantification of four independent experiments as in **c**. Data are presented as mean ± SEM (n=4). n.s. not significant, *** *p* <0.001, **** *p* <0.0001 using two-way ANOVA with Holm-Sidaks multiple comparison tests. **e** Representative images of COS-7 cells expressing FLAG-tagged CNAβ2, CNAβ1 or CNAβ1 double mutant C483/C493S with the PM marker Venus-tagged Rit (green). Fixed cells immunostained with anti-FLAG (red) and anti-GM130 (blue). Scale bar = 15 µm. **f** Graph on top: Co-localization of FLAG signal (as in e) with Golgi marker GM130. Data are presented as median Pearson’s coefficients. At least 100 cells were analyzed across three independent replicates. n.s not significant, Asterisks denote statistical significance: **** *p* <0.0001, using one-way ANOVA followed by Kruskal-Wallis test. **e** Graph at the bottom: PM localization is quantified as the anti-FLAG signal intensity at the cell periphery (defined in Supplementary Fig. 1d) over total cell intensity. Refer to methods for details. Data are presented as median. At least 70 cells were imaged across three independent replicates. **** *p* <0.0001 using one-way ANOVA followed by Kruskal-Wallis test. **g** Representative immunoblot of Acyl-PEG exchange assay done in COS-7 cells transfected with FLAG-CNAβ2, FLAG -CNAβ1 (WT or cysteine mutants: singles and double) or EFR3B-FLAG. The number of PEGylation (reflecting S-palmitoylation) events are indicated by asterisks (*). Arrowhead indicates non-specific antibody band. n ≥3 experiments for all constructs

To date, efforts to discover CN-regulated processes have focused on canonical CN isoforms, leaving CNAβ1 significantly understudied. Interestingly, the few published studies about this isoform demonstrate that it has unique physiological roles. For example, CNAβ1 overexpression in mouse cardiomyocytes is cardio-protective following myocardial infarction, rather than pro-hypertrophic, as observed for canonical CNAβ2 ^22–24^. Furthermore, mice specifically lacking CNAβ1 are viable, but develop cardiac hypertrophy and exhibit metabolic alterations ^24^. CNAβ1 also regulates the differentiation of mouse embryonic stem cells and activates mTORC2/AKT signaling through an undetermined mechanism that may be independent of its catalytic activity ^14, 22, 25^. Additionally, unlike CNAβ2, CNAβ1 does not dephosphorylate NFAT ^22^, and its direct substrates are yet to be identified. Thus, elucidation of these targets promises to reveal novel aspects of Ca^2+^ and CN signaling.

Some of the best-characterized pathways that generate intracellular Ca^2+^ signals are initiated by ligand binding to Gq-protein coupled receptors (GPCR), causing phospholipase C (PLC) to hydrolyze phosphatidylinositol 4,5-biphosphate [PI(4,5)P_2_ or PIP_2_] into diacylglycerol (DAG) and inositol triphosphate (IP_3_). These products activate protein kinase C (PKC) and intracellular Ca^2+^ release, respectively ^26^. Therefore, sustained Ca^2+^ signaling through GPCRs requires continued phosphorylation of plasma membrane (PM) phosphatidylinositol (PI) to generate phosphatidylinositol 4-phosphate (PI4P), the precursor of PI(4,5)P_2_. Indeed, studies monitoring the PM phospholipid levels in real-time reveal that during GPCR signaling, concomitant with PIP2 depletion, PI4P synthesis increases through the activity of phosphatidylinositol 4-kinase IIIα (PI4KA) ^27, 28^.

PI4KA is recruited to the PM by associating with at least two accessory proteins, EFR3A/B and TTC7A/B, a mechanism that is conserved from yeast to mammals ^29–32^. EFR3, which is stably associated with the PM due to its palmitoylation, serves as the membrane anchor for this complex ^33^. TTC7 (Ypp1 in yeast) binds to both EFR3 and PI4KA (Stt4 in yeast) and acts as the shuttle. A third protein, either FAM126A (Hyccin) or FAM126B, present only in higher eukaryotes, is an essential, regulatory component that stabilizes the TTC7-PI4KA interaction in the cytosol, and enhances recruitment of PI4KA to the PM ^30^. Recent structural studies show that PI4KA/TTC7/FAM126A heterotrimers form a dimer, and this super-assembly likely stabilizes and orients the PI4KA active site toward the membrane to promote its activity ^34^. Furthermore, a recent biochemical study reveals that the disordered C-terminus of FAM126A, which is not present in existing structural data, modulates the PI4KA catalytic activity *in vitro* through an unknown mechanism ^35^. This intricate structure suggests that both the assembly and activity of the PI4KA complex are tightly regulated. In yeast, PI4KA recruitment to the PM is regulated by phosphorylation ^32^. However, in mammals, how the assembly and/or activity of the PI4KA complex is regulated remains to be elucidated.

In this work, we discover novel functions for CN by focusing on the CNAβ1/CNB isozyme. We demonstrate that unlike the cytosolic canonical CNAβ2, CNAβ1 localizes to cellular membranes, primarily to the PM and Golgi apparatus, via palmitoylation of two conserved cysteines within its unique C-terminus. Palmitoylation of CNAβ1 is dynamic and is reversed by the ABHD17A thioesterase leading to its redistribution and suggesting that dynamic palmitoylation regulates CNAβ1 signaling *in vivo*. To identify potential CNAβ1 substrates we carried out affinity purification coupled to mass spectrometry (AP-MS) which revealed CNAβ1-specific interactors to be largely membrane-associated, and unexpectedly identified all four members of the PI4KA complex. Using *in vivo* and *in vitro* analyses, including hydrogen deuterium exchange mass spectrometry (HDX-MS), we identified multiple sites of CN-PI4KA complex association, including a direct interaction with a short linear motif, PSISIT, within the unstructured C-terminal tail of FAM126A. Our studies establish FAM126A as a CN substrate that preferentially interacts with CNAβ1 at the PM. Finally, we uncover a role for CN in the production of PI4P at the PM by PI4KA in response to ligand induced signaling from the type-3 muscarinic receptor. In total, this work discovers a new CN-regulated signaling pathway that highlights the PI4KA complex as a regulatory target and demonstrates that palmitoylation dictates substrate specificity of the non-canonical CNAβ1 isoform.

## Results

### CNAβ1 localizes to the plasma membrane, Golgi apparatus and intracellular vesicles

We sought to investigate the unique functions of CNAβ1 by characterizing its *in vivo* properties. First, we analyzed the intracellular distribution of CNAβ1, which was previously found to be Golgi-associated in mouse embryonic stem cells ^25^. Using subcellular fractionation of COS-7 cells expressing FLAG-tagged CNAβ1 or CNAβ2, we confirmed that CNAβ1 was highly enriched in the membrane fraction while CNAβ2 was primarily found in the cytosol fraction (Fig. 1 c-d). Furthermore, indirect immunofluorescence of these cells revealed that CNAβ1 localized to the PM, where it overlapped significantly with a co-expressed PM marker (Venus-RIT) ^36^, the Golgi apparatus, where it co-localized with the Golgi protein GM130, and to intracellular vesicles. By contrast, CNAβ2 was predominantly present in the cytoplasm with minimal co-localization with either membrane marker (Fig. 1 e-f). Similar observations were also made in HeLa cells (Supplementary Fig.1a).

### CNAβ1 is palmitoylated at two conserved cysteines unique to its C-terminal tail

S-Palmitoylation, the reversible addition of a 16-carbon fatty acid chain to cysteine residues via a thioester linkage, allows proteins lacking a transmembrane domain to associate with cellular membranes. The alternative C-terminus of CNAβ1 contains two highly conserved cysteines, C483 and C493 (Fig. 1a and b), one of which (C493) is predicted as a high-confidence S-palmitoylation site ^37^. We reasoned that this modification might mediate CNAβ1 membrane association, particularly as C483 is contained within a previously defined “Golgi localization domain” ^25, 38^. First, we investigated this possibility using acyl resin-assisted capture (Acyl-RAC), during which the thioester linkage in palmitoylated cysteines is cleaved with hydroxylamine (NH_2_OH) to allow protein binding to thiopropyl-sepharose beads ^39^. Supporting our hypothesis, Acyl-RAC analysis of FLAG-CNAβ2, FLAG-CNAβ1 or EFR3B-FLAG expressed in COS-7 cells revealed the presence of S-palmitoylated cysteines in EFR3B and CNAβ1, but not CNAβ2 (Supplementary Fig. 1b). CNAβ1 mutants containing either single or double serine substitutions at C483 and/or C493, from here on referred to as CNAβ1^C483S^, CNAβ1^C493S^ and CNAβ1^C2S^ respectively, were not captured by Acyl-RAC suggesting that at least one of these residues is palmitoylated (Supplementary Fig. 1b). To further determine the stoichiometry of CNAβ1 palmitoylation, we used acyl-PEG exchange (APE) in which the palmitate groups on modified cysteines are removed by hydroxylamine and replaced with a mass-tag (mPEG) that causes a 5 kDa mass-shift for each acylated cysteine ^40^. For these experiments we used two controls: EFR3B which contains three palmitoylated cysteine residues and the ER chaperone calnexin with two sites of palmitoylation ^33, 41^. As expected, EFR3B-FLAG showed three distinct bands corresponding to mass-shifts and the endogenous calnexin showed two mass-shifts in all samples (Fig. 1g). As expected, the cytosolic FLAG-CNAβ2 showed no shifts. FLAG-CNAβ1, however, displayed two distinct mass-shifted forms indicating two sites of palmitoylation. Interestingly, no changes in electrophoretic mobility were observed for CNAβ1^C483S^, CNAβ1^C493S^ or CNAβ1^C2S^, indicating that both cysteines are required for stable palmitoylation. Thus, we infer that CNAβ1 palmitoylation on both sites is a cooperative process, similar to that described for calnexin ^42^.

### Palmitoylation is required for CNAβ1 membrane association

To further investigate a role for palmitoylation in the membrane association of CNAβ1, we first metabolically labelled COS-7 cells with the palmitate analog, 17-octadecynoic acid (17-ODYA) and showed that upon subcellular fractionation, the majority of the 17-ODYA-labelled CNAβ1 was in the membrane fraction (Supplementary Fig. 1c). Next, we analyzed the fractionation of palmitoylation-defective mutants CNAβ1^C483S^, CNAβ1^C493S^ and CNAβ1^C2S^ which, in contrast to wildtype CNAβ1, were predominantly enriched in the cytosolic fraction (Fig. 1c, d). Finally, we examined the localization of each mutant using indirect immunofluorescence. As expected, FLAG-CNAβ1^C493S^ and FLAG-CNAβ1^C2S^ mutants were cytosolic and did not co-localize with either PM or Golgi membrane markers (Venus RIT and GM130, respectively) (Fig. 1e, f and Supplementary Fig. 1e, f). Interestingly, although FLAG-CNAβ1^C483S^ was predominantly cytosolic, a minority of cells exhibited weak Golgi and PM localization (Supplementary Fig. 1e (top panel, red and green boxes), f), suggesting that this mutant might be palmitoylated at low levels that is insufficient for stable membrane association. Thus, we speculate that Cys493 may be the priming palmitoylation site that promotes efficient palmitoylation of Cys483. In sum, these analyses show that both Cys483 and Cys493 are palmitoylated and that dual palmitoylation is required for the stable association of CNAβ1 with membranes, particularly with the PM. Therefore, the unique lipidated C-terminal tail of CNAβ1 confers distinct localization to this isoform.

### CNAβ1 palmitoylation is dynamically regulated

Protein palmitoylation is reversed by acyl protein thioesterases (depalmitoylases) and dynamic palmitoylation regulates the localization and function of many signaling proteins including RAS GTPases ^43^. To determine if palmitates on CNAβ1 actively turn over in cells, we performed a pulse-chase experiment by briefly labelling COS-7 cells expressing FLAG-CNAβ1 with 17-ODYA and the methionine analog L-azidohomoalanine (L-AHA), followed by immunopurification and visualization of CNAβ1, using dual-click chemistry to determine levels of 17-ODYA and L-AHA incorporation over time (Fig. 2a). In cells treated with Palmostatin B (Palm B), a pan inhibitor of depalmitoylases, the ratio of 17-ODYA/L-AHA incorporation into FLAG-CNAβ1 increased over time relative to the control (DMSO treated) cells, establishing that palmitoylation of CNAβ1 is reversed by endogenous depalmitoylases (Fig. 2b, c).

**Figure 2:**
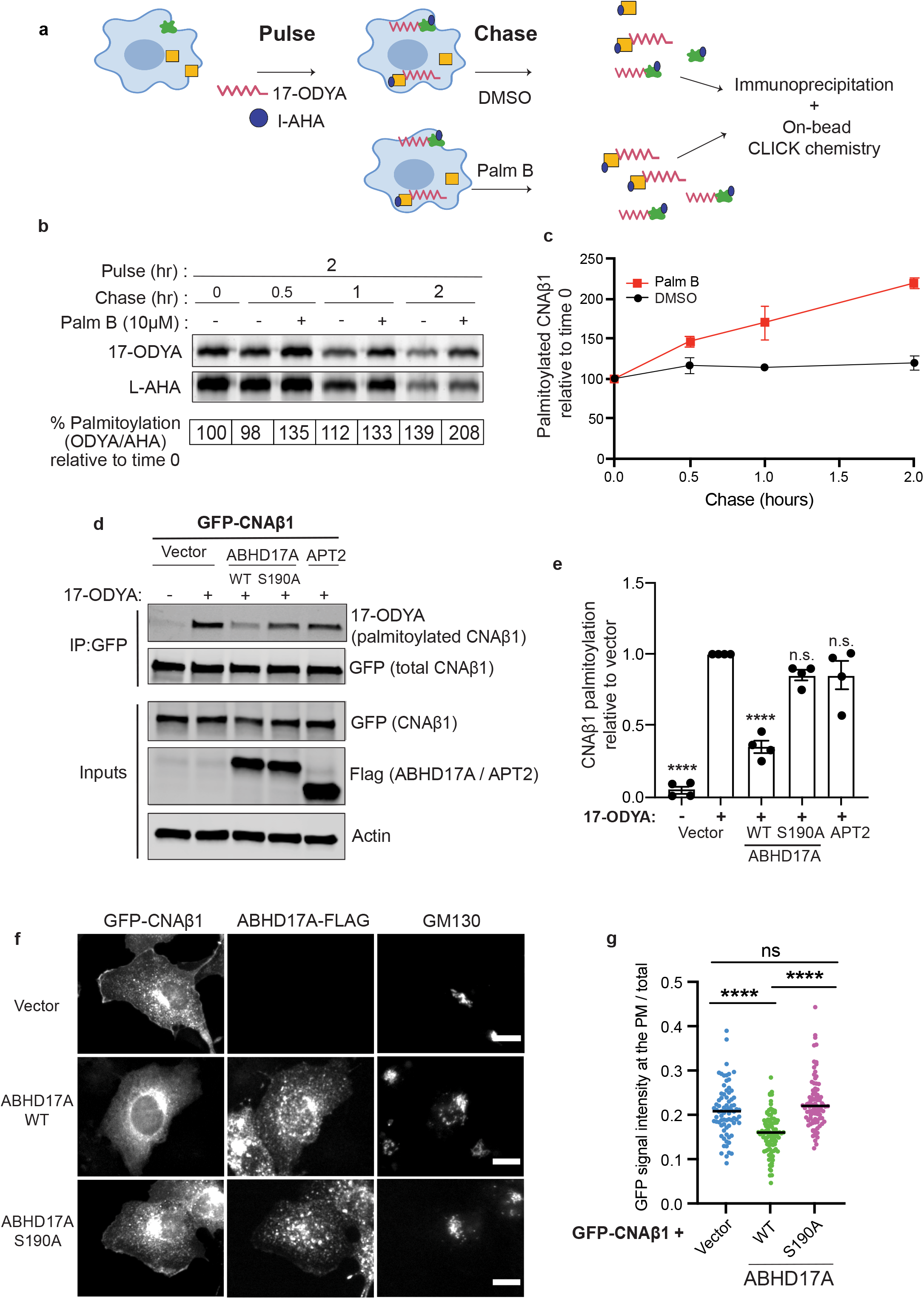
CNAβ1 palmitoylation is dynamic: ABHD17A expression promotes CNAβ1 depalmitoylation and alters CNAβ1 subcellular localization. **a** Schematic diagram of the pulse-chase experiment using analogs of palmitate (17-ODYA) and methionine (L-AHA) coupled to CLICK chemistry used in this study. **b** Pulse-chase analysis of palmitate turnover on FLAG-CNAβ1 by dual-click chemistry as described in **a** in the presence of DMSO or pan-depalmitoylase inhibitor Palm B. Representative in-gel fluorescence scans showing dual detection of 17-ODYA and L-AHA using Alexa Fluor 647 and Alexa Fluor 488, respectively. **c** Time course of FLAG-CNAβ1 depalmitoylation in DMSO- and Palm B-treated cells after normalizing 17-ODYA to L-AHA signals at each chase time. Data are mean ± SEM, n= 2 **d** Analysis of GFP-CNAβ1 palmitoylation co-expressed with vector, ABHD17A-FLAG (WT or S190A) or FLAG-APT2, using metabolic labelling with 17-ODYA. Representative immunoblot illustrates total CNAβ1 using anti-GFP and 17-ODYA detected using streptavidin following CLICK chemistry with azide-Biotin. Anti-FLAG shows amount of ABHD17A and APT2 expression. **e** Level of GFP-CNAβ1 palmitoylation in **d** is quantified by the streptavidin signal (17-ODYA) / total protein signal (GFP) and normalized to vector control. Data are mean ± SEM (n=4). n.s. not significant, **** *p* <0.0001 using one-way ANOVA with Dunnett’s multiple comparison tests. **f** Representative IF images of fixed, COS-7 cells co-expressing GFP-CNAβ1 with vector, ABHD17A-FLAG (WT or S190A). Scale bar = 15 µm. **g** Images in **f** quantified, scatter plot represents the intensity of GFP-CNAβ1 at the PM relative to total GFP signal. At least 75 cells quantified per condition across three replicates. n.s. not significant, **** *p* <0.0001 using one-way ANOVA followed by Kruskal-Wallis test.

In mammals, two classes of thioesterases are responsible for the removal of palmitate groups resulting in protein depalmitoylation. The soluble, acyl protein thioesterases (APT1, APT2) and the membrane associated α/β hydrolase domain proteins (ABHDs) display substrate specificity likely due to their differential subcellular distribution ^43–45^. Among the ABHD family, we focused on ABHD17A for which the catalytic properties and intracellular localization to the PM, Golgi and endosomes are well-characterized ^44^. To identify which of these thioesterases regulate the palmitate turnover in CNAβ1, we carried out metabolic labeling with 17-ODYA in COS-7 cells expressing GFP-CNAβ1 together with a vector, ABHD17A-FLAG (WT or the catalytically impaired mutant S190A ^44^), FLAG-APT2 or mCherry-APT1. Overexpression of ABDH17A WT, but not S190A, dramatically decreased the 17-ODYA labelling of GFP-CNAβ1 (Fig. 2d, e), and resulted in redistribution of GFP-CNAβ1 from the PM to the cytosol and Golgi (Fig. 2f, g). In contrast, overexpression of APT2 (Fig. 2d, e) or APT1 (Supplementary Fig. 2a, b) did not alter palmitoylation of CNAβ1 suggesting that CNAβ1 is depalmitoylated specifically by ABHD17A. Together, these data reveal that palmitoylation of CNAβ1 is dynamic, which provides a potential mechanism to regulate its localization *in vivo*. In sum, our findings demonstrate that CNAβ1 has cellular properties distinct from canonical CN isoforms, which led us to investigate whether this isoform has specific substrates and functions at membranes.

### Affinity purification and mass spectrometry identifies CNAβ1-specific interactors

Previous studies report that CNAβ1, unlike canonical CN isoforms, does not activate or interact with NFAT ^22^. Indeed, when FLAG-NFATC1 was co-expressed in HEK293 T-REx cell lines that inducibly express GFP, GFP-CNAα, GFP-CNAβ2 or GFP-CNAβ1, immunoprecipitation using anti-GFP antibody confirmed that NFATC1 co-purifies with CNAα and CNAβ2, but not with CNAβ1. By contrast, the CNB regulatory subunit, was recovered to the same extent with all three CN isoforms (Supplementary Fig. 3b). Next, to identify CNAβ1-specific interactors which might include substrates, we turned to affinity purification coupled to mass spectrometry (AP-MS). HEK293 T-REx cell lines were developed that express either 3X FLAG-tagged-GFP, -CNAβtrunc lacking the C-terminal tail (aa 1-423; truncated after calmodulin binding site), canonical -CNAβ2, or -CNAβ1 (Fig. 3a). Following immunoprecipitation, co-purifying proteins were identified using label-free, quantitative mass spectrometry and the 3X FLAG-tagged-GFP control was used to eliminate non-specific interactors. In total, 51 high confidence CN-interacting proteins (defined as those with a bayesian false discovery rate (BFDR) ≤ 1%) were identified (Supplementary Fig. 3a, Supplementary Table 1). As expected, some established CNA interactors, including the CNB subunit (PPP3R1) and the inhibitor RCAN3, were identified with all CNAβ constructs (Fig. 3b, Supplementary Fig. 3a). Of these 51 proteins, 12 were previously identified as CN-interactors and several, including BRUCE, FAM126A and GSK3β contain predicted CN binding motifs (LxVP or PxIxIT) confirming the validity of our data set ^6, 7, 46^.

**Figure 3:**
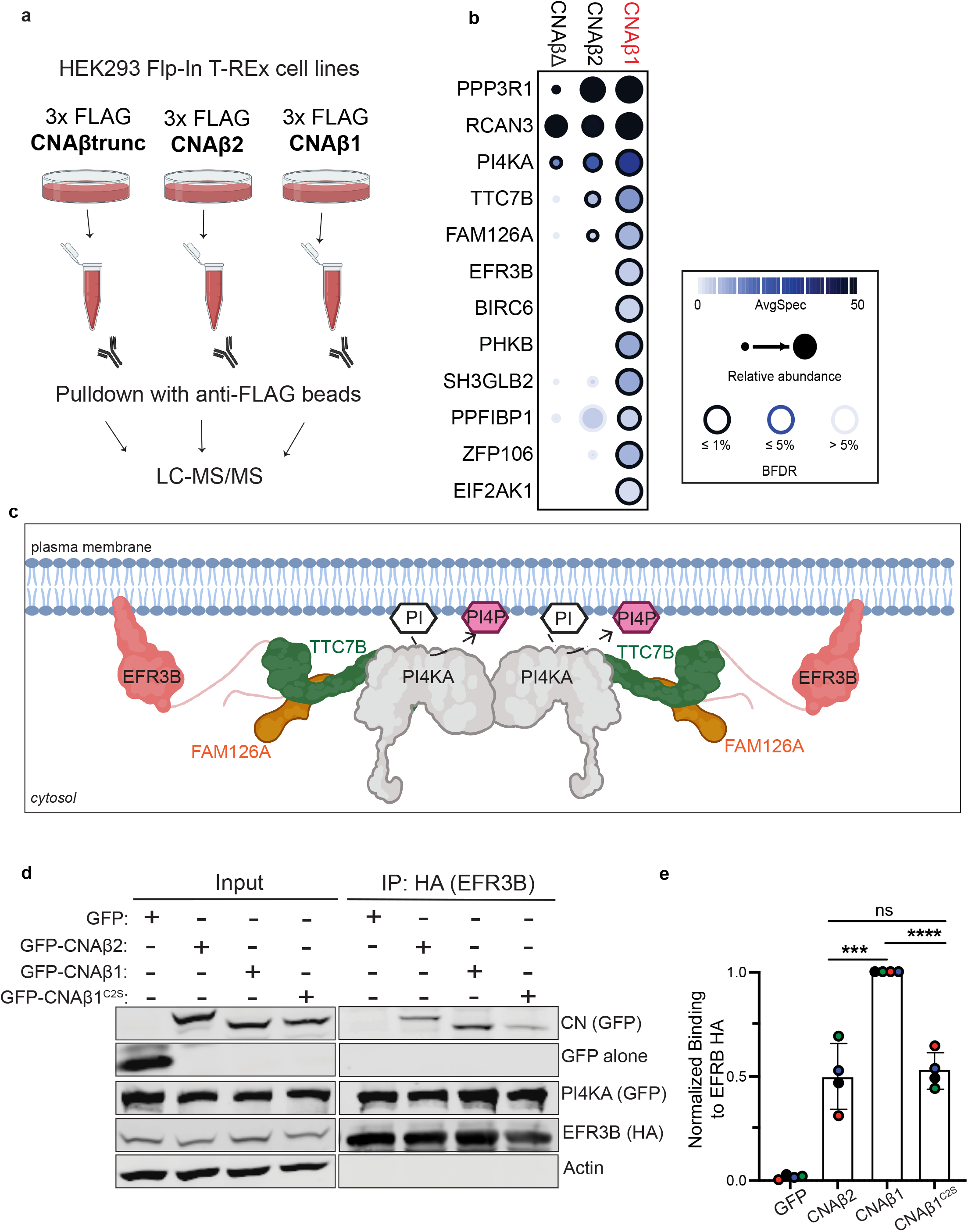
CNAβ1-enriched interactors are membrane-associated. **a** Schematic of the experimental plan used for AP-MS analyses. **b** Dotplot of AP-MS results including CNAβ1-enriched interactors (those with spectral counts ≥ 1.5x more for CNAβ1 than other baits). Node edge color corresponds to bayesian false discovery rate (BFDR), node size displays prey abundance and node darkness represents number of spectral counts. Full results reported in Supplementary Fig. 3a and Supplementary Table 1. **c** Cartoon representation of the structural organization of the phosphatidylinositol 4-kinase complex that comprises PI4KA (grey), FAM126A (orange), TTC7B (green) and EFR3B (pink). Phosphatidylinositol, PI (white); phosphatidylinositol 4-phosphate, PI4P (purple). **d** Immunoblot analysis of the anti-GFP immunoprecipitates from inducible Flp-In-Trex cells expressing GFP-CNAβ2, CNAβ1 or CNAβ1^C2S^ (C483S/C493S), transfected with EFR3B-HA, TTC7B-MYC, FLAG-FAM126A and GFP-PI4KA. **e** Amount of GFP-CNAβ2 and GFP-CNAβ1^C2S^ co-purified with EFR3B-HA, quantified as bound GFP signal/ bound HA signal normalized to input. Data are the mean ± SD (n=4). n.s. not significant, *** *p* < 0.001, **** *p* <0.0001 by unpaired-t-test.

Excitingly, several proteins were enriched in immunoprecipitates of CNAβ1 relative to CNAβ2 or CNAβtrunc (Fig. 3b). Consistent with the intracellular distribution of CNAβ1, the majority of these were membrane-associated, including Baculoviral IAP repeat-containing protein 6 (BIRC6/BRUCE), which localizes to the Golgi and endosomes ^47^, PM-associated Phosphorylase B kinase regulatory subunit (PHKB), cell junction protein Liprin-Beta 1 (PPFIBP1) ^48^, and endosomal SH3 and BAR domain-containing protein endophilin B2 (SH3GLB2) ^49^. And most strikingly, all subunits of the large PM-associated PI4KA complex, composed of EFR3B, FAM126A (Hyccin), TTC7B and PI4KA (PI4KIIIα), were identified (Fig. 3b, c) ^29, 30, 32^. Together, these findings suggest that CNAβ1 interacts with a unique set of membrane-associated proteins which may represent novel CNAβ1-regulated substrates and pathways.

### CNAβ1 interacts with the PI4KA complex at the plasma membrane

We focused our attention on the PI4KA complex, which is endogenously expressed at very low levels. To ensure the balanced expression of PI4KA complex components in our studies, we engineered a single plasmid that harbors the DNA sequences encoding EFR3B, TTC7B and FAM126A separated by the viral 2A linkers T2A and P2A, respectively (Supplementary Fig. 3d). During translation, 2A peptides are cleaved leading to the expression of each protein separately in constant stoichiometry ^50^. Efficient cleavage of this plasmid and the proper expression of each component was verified in both HeLa (Supplementary Fig. 3f) and HEK293 cells (data not shown). Immunofluorescence analyses of HeLa cells further confirmed the expected PM localizations for EFR3B, TTC7B, FAM126A (Supplementary Fig. 3e) and co-expressed PI4KA (Supplementary Fig. 5c), indicating that the complex was functional. Using this expression system, we first performed reciprocal immunoprecipitation experiments to validate the enriched interaction of the PI4KA complex with CNAβ1 compared to the canonical CNAβ2. HEK293 T-REx cells inducibly expressing GFP-FLAG control, GFP-CNAβ2 or GFP-CNAβ1 were transfected with the EFR3B-HA, TTC7B-MYC, FLAG-FAM126A-containing plasmid together with GFP-PI4KA. EFR3B-HA was immunoprecipitated from cell lysates and the co-purifying proteins were analyzed. As expected, GFP-PI4KA efficiently co-purified with EFR3B indicating functional complex formation. Supporting our AP-MS results, association of GFP-CNAβ1 with EFR3B was significantly higher than that of either GFP-CNAβ2 or the palmitoylation defective GFP-CNAβ1^C2S^ (Fig. 3d, e). Thus, CNAβ1 preferentially interacts with the PI4KA complex due to its unique PM localization, which is mediated by palmitoylation.

### FAM126A has a putative PxIxIT motif that mediates binding to CN

Computational predictions identified a highly conserved sequence motif within the intrinsically disordered C-terminal tail of FAM126A, ^512^PSISIT^517^, matching the consensus of the PxIxIT motif that mediates binding to CN ^6, 7^ (Fig. 4a, Supplementary Fig. 4a). To determine whether this sequence binds to CN, we first fused a 16-mer peptide containing this sequence from FAM126A to GST and tested its co-purification with the recombinant, HIS-tagged CN heterodimer *in vitro*. GST fused to the PxIxIT from NFATC1 was used as a positive control. As expected, the FAM126A peptide efficiently co-purified with wildtype HIS-CN but not with mutant CN (NIR) which is defective for PxIxIT-docking ^51^. Mutating key residues of the PSISIT sequence to alanine (FAM126A^ASASAA^), also disrupted the interaction with CN showing that this sequence mediates direct binding to CN *in vitro* (Supplementary Fig. 4b, c).

**Figure 4:**
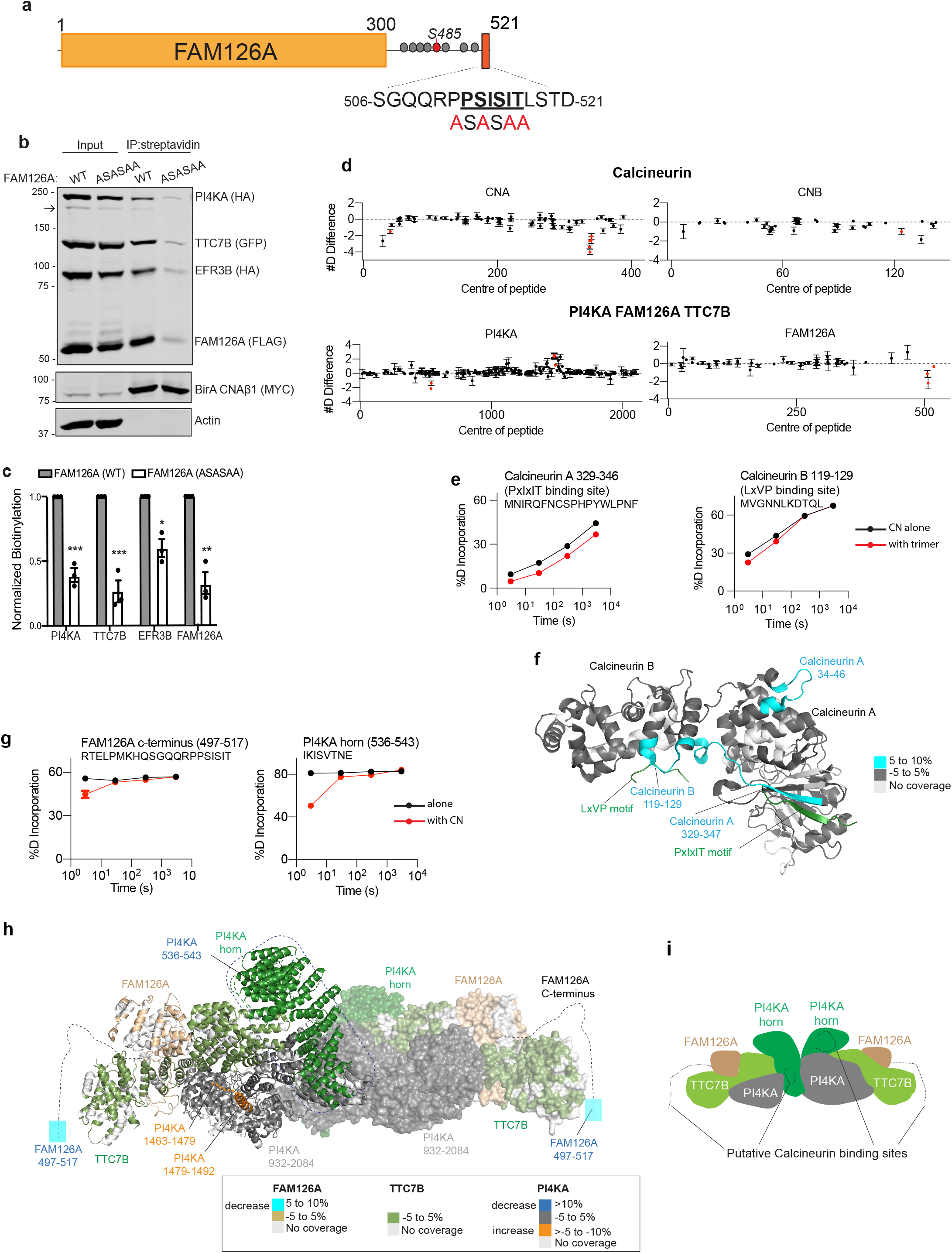
FAM126A has a CN binding PxIxIT motif. **a** Schematic of FAM126A with the 16-mer peptide (aa.506-521) containing the PSISIT sequence bolded and underlined. ASASAA mutations are shown in red. Gray circles denote known phosphorylation sites. The red circle denotes phospho-Ser 485. **b** Representative immunoblot showing the biotinylation of the expressed PI4KA complex containing FLAG-FAM126A (WT or ASASAA) by MYC-BirA*-CNAβ1 in HeLa cells. The arrow shows a small fraction of the uncut P2A protein. **c** Biotinylation quantified as bound signal/ MYC bound signal normalized to input signal/Actin signal. Data are the mean ± SEM (n=3). n.s. not significant, * *p*<0.05, ** *p*< 0.01, *** *p*<0.001 using unpaired t-tests. **d-e** HDX data for CN heterodimer (CNA/CNB) and the PI4KA/TTC7B/FAM126A trimer. **d** Differences in the number of deuteron incorporation for all analyzed peptides over the deuterium exchange (HDX) time course. Proteins with significant differences between apo and complex state are shown. Every point represents the central residue of a peptide. Peptides with significant HDX (>5%, 0.5 Da, and an unpaired t-test *p*<0.01) are highlighted in red. **e** Deuterium incorporation differences between selected CNA and CNB peptides in the presence of PI4KA/TTC7B/FAM126A trimer are shown. Black lines represent CN in its apo state, red lines represent CN co-incubated with the PI4KA trimer. **f** Maximum significant differences in HDX observed in CNA/CNB across all time points upon incubation with PI4KA trimer are mapped onto the structure of CN ^58^ (PDB: 6NUC). Peptides are colored according to the legend. The PxIxIT and LxVP motifs identified in NHE1^58^ are highlighted in green. **g** Deuterium incorporation differences in selected FAM126A and PI4KA peptides displaying significant decreases in amide exchange in the presence of CN are shown. All error bars show the S.D. (n = 3). **h** Maximum significant differences in HDX observed for PI4KA/FAM126A/TTC7B trimer in the presence of CN are mapped onto the structure of the PI4KA trimer ^34^ (PDB:6BQ1). Dotted lines depict the unresolved regions in the PI4KA/TTC7/FAM126A structure, colors represent the percentages of significant differences in exchange. **i** Schematic of the PI4KA complex with putative CN interaction sites. The full set of peptides and source data can be found in Supplementary Fig. 4d-e and Supplementary Table 3, respectively.

Next, to investigate whether FAM126A-CNAβ1 association is PxIxIT-dependent *in vivo,* we used proximity-dependent labeling (BioID) with the promiscuous biotin ligase, BirA*, which sensitively detects the low affinity interactions seen between CN and substrates ^6, 52, 53^. We transfected HeLa cells expressing BirA-fused CNAβ1 with HA-PI4KA and the EFR3B-HA, TTC7B-MYC, FLAG-FAM126A-containing plasmid described above with either FAM126A ^WT^ or CN-binding-defective FAM126A^ASASAA^. Consistent with AP-MS results, each component of the PI4KA complex was biotinylated by BirA-fused CNAβ1 (Fig. 4b, c) and as expected, FAM126A^WT^ was significantly more biotinylated than FAM126A^ASASAA^ (Fig. 4b, c). Interestingly, biotinylation of other complex members, i.e, TTC7B, PI4KA and EFR3B was also reduced in the presence of FAM126A^ASASAA^. In sum, these findings identify PSISIT as a direct a CN-binding motif in FAM26A and suggest that this sequence promotes interaction of CNAβ1 with the entire PI4KA complex.

### Hydrogen/deuterium exchange maps CN-PI4KA complex interaction sites

The cryo-EM structure of PI4KA-TTC7B-FAM126A fails to resolve the unstructured, disordered C-terminal tail of FAM126A, which contains the CN-binding motif ^34^. Therefore, we turned to hydrogen deuterium exchange coupled to mass spectrometry (HDX-MS) to map the CN-PI4KA complex interaction and identify any conformational changes that occur upon binding ^35, 54^. HDX-MS measures the exchange rate of amide hydrogens with deuterium-containing buffer, which acts as a sensitive probe of secondary structure dynamics ^55^. The CNA/CNB heterodimer was produced in *Escherichia coli* and recombinant PI4KA in complex with TTC7B and FAM126A was purified from insect cells as previously described ^35^. The PI4KA/TTC7B/FAM126A trimer and CNA/CNB were exposed to pulses of deuterium when incubated alone or together, with CN in excess over the PI4KA trimer (Fig. 4d). Localization of differences in H/D exchange requires proteolysis into peptides, with sequence coverage for PI4KA, FAM126A, TTC7B and Calcineurin A (catalytic) and B (regulatory) subunits of 77.6%, 80.9%, 84.2%, 89% and 89.3%, respectively (Supplementary Table 2). Following addition of deuterium-containing buffer (D_2_O), reactions were quenched at indicated times (3s, 30s, 300s, 3000s) and the resulting shifts in mass upon deuterium-incorporation was analyzed with mass spectrometry. Peptides that showed differences in amide exchange greater than 0.5Da and 5% at any time point and had unpaired t-test values of *p* <0.01, across three replicates, were considered significant.

Co-incubation of CNA/CNB with the PI4KA/TTC7B/FAM126A trimer resulted in a large decrease in hydrogen-deuterium exchange in the well-characterized PxIxIT docking groove in CNA (aa 329-346) (Fig. 4e, f), consistent with the PxIxIT-mediated interaction we demonstrated between FAM126A and CN. We also observed decreased amide exchange in the N-terminus of CNA (aa 34-36) suggesting that previously unidentified conformational changes occur upon substrate binding (Fig. 4d, f and Supplementary Fig. 4d). Interestingly, a region in CNB that forms part of the LxVP-binding groove (aa 119-129) also showed significantly decreased amide incorporation which may indicate that additional, as yet unidentified, LxVP-mediated interactions occur between the PI4KA trimer and CN (Fig. 4d, e and f). As for the PI4KA complex, while no significant changes in amide exchange were seen in TTC7B (Supplementary Fig. 4e), a few regions in both FAM126A and PI4KA showed significant changes in deuterium exchange in the presence of CNA/CNB. In FAM126A, exchange decreased significantly in the region within the C-terminal tail that contains the PSISIT sequence (aa 497-517) consistent with CN-binding to this site (Fig. 4d, g and h). In PI4KA an unstructured region within the α-solenoid domain (aka the horn) (aa 536-543) showed a decrease in deuterium incorporation (Fig. 4d, g and h), indicating the formation of secondary structure either due to direct interaction with CN or as an indirect consequence of CN binding to FAM126A. Interestingly, this region contains a PxIxIT-like peptide sequence, “IKISVT”, which may be a novel, non-canonical CN binding motif. In addition, a set of peptides identified in PI4KA between residues 1463-1492 showed increased amide exchange (Fig. 4h, Supplementary Fig. 4d, e) revealing a conformational change that occurs in the presence of CN. Overall, these studies indicate multiple sites of contact between CN and PI4KA/TTC7B/FAM126A trimer suggestive of a regulatory interaction.

### FAM126A is a novel CN substrate

To examine whether CN regulates phosphorylation of the PI4KA complex we focused on FAM126A because of its small size (∼58 kDa), the presence of a confirmed CN-binding motif and several identified sites of phosphorylation ^56^. First, we expressed FLAG-FAM126A^WT^ or CN-binding defective FAM126A^ASASAA^, alone or together with TTC7B-MYC and EFR3B-HA and examined their electrophoretic mobility via SDS-PAGE and immunoblot analysis. Slower migrating forms of FAM126A were observed that were enhanced in FAM126A^ASASAA^ compared to FAM126A^WT^ (labelled PI and PII in Fig. 5a, lane 2 vs 5). Notably, these shifts were present only when FAM126A was co-expressed with the other components (Fig. 5a, lane 1 vs 2 or lane 4 vs 5), especially EFR3B, the membrane anchor for the complex ^30^ (Supplementary Fig. 5a). These slower migrating forms, indicative of hyperphosphorylation, suggest that that FAM126A is phosphorylated only when associated with the PM-localized PI4KA complex, and that CN dephosphorylates FAM126A in a PxIxIT-dependent manner. To further analyze FAM126A phospho-regulation, we mutated several serine and threonine residues observed to be phosphorylated ^56^ to the non-phosphorylatable amino acid alanine. Remarkably, mutating serine 485 (FAM126A^S485A^) altered mobility shifts in FAM126A, eliminating PII and reducing PI (Fig. 5a, lane 2 vs 3 and lane 5 vs 6). This suggests that Ser485 is one target of phosphorylation and that additional sites likely contribute to the observed shifts. To analyze the phosphorylation status of Ser485 in FAM126A, we generated a phospho-specific antibody for this site (anti-pFAM126A S485). The specificity of this antibody is demonstrated by analyses of HeLa cells expressing FAM126A mutants (S485A, ASASAA or ASASAA+S485A) with or without EFR3B and TTC7B co-expression, where this antibody specifically recognized both slower-migrating FAM126A forms (PI and PII, Fig. 5a). Notably, no signal was detected for FAM126A^S485A^ or when FAM126A was expressed alone, and the total signal was significantly higher for FAM126A^ASASAA^ compared to FAM126A^WT^. Moreover, indirect immunofluorescence using anti-pFAM126A S485 antibody showed enriched signal at the PM, further indicating that FAM126A is phosphorylated when the PI4KA complex is associated with the PM (Supplementary Fig. 5b).

**Figure 5:**
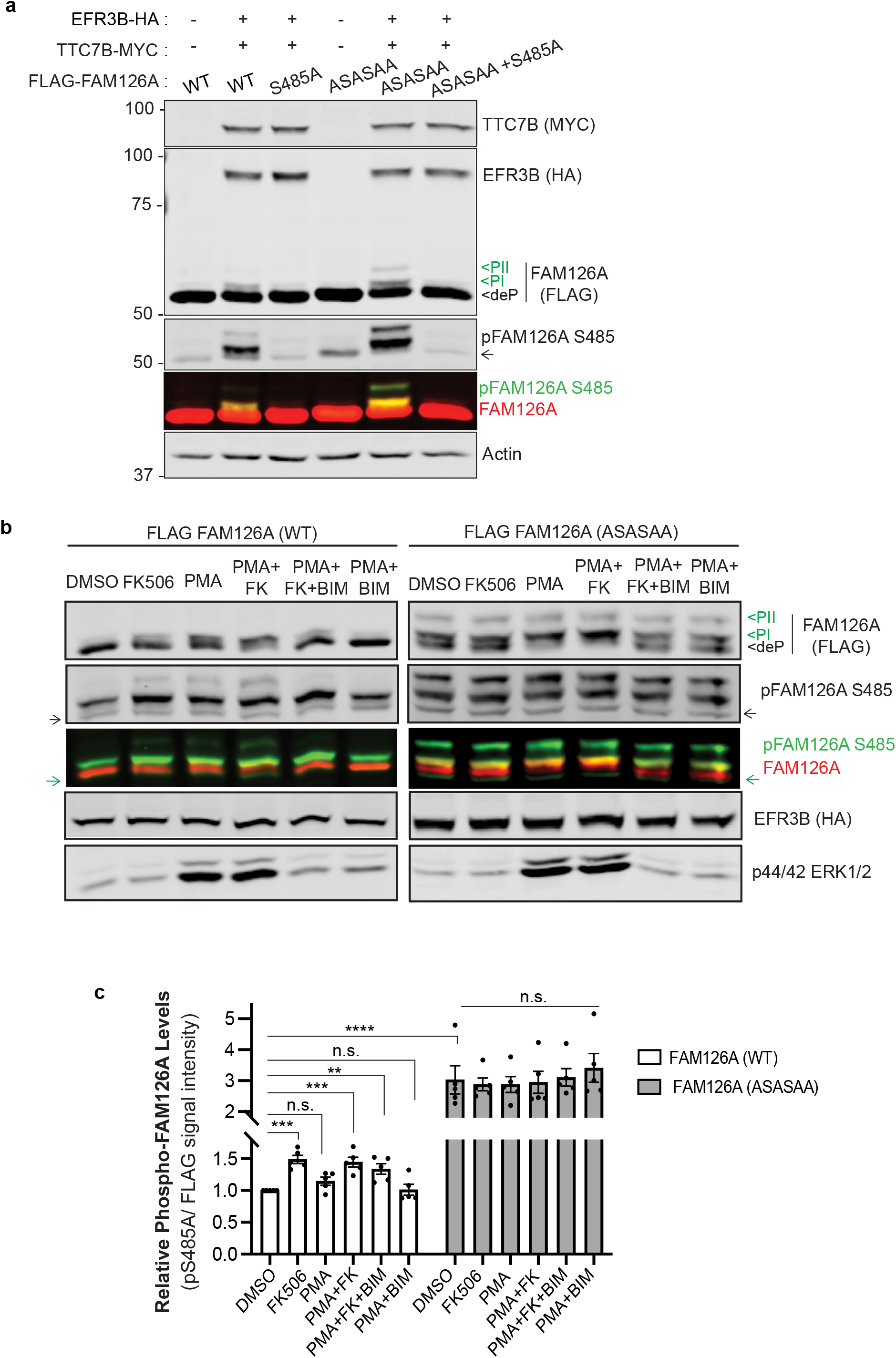
FAM126A is a novel CN substrate. **a**. Representative immunoblot showing the electro-mobility shifts observed for FLAG-FAM126A when expressed in HeLa cells. Lysates expressing FLAG-FAM126A (WT, S485A, ASASAA or ASASAA+S485A) in the presence or absence of EFR3B-HA/ TTC7B-MYC are analyzed using MYC, HA, FLAG (red bands) and a phospho-specific pFAM126A S485A (green bands) antibodies. PI and PII: phosphorylated states; deP: dephoshorylated state. **b** Immunoblot analysis of FLAG FAM126A (WT or ASASAA) phosphorylation status in HeLa cells co-expressing FLAG-FAM126A, TTC7B-MYC and EFR3B-HA across various treatments: DMSO (vehicle), FK506 (CN-inhibitor), PMA (PKC activator), BIM (PKC). Lysates were resolved by SDS-PAGE and analyzed using anti-FLAG (red), anti-HA and anti-pFAM126A S485A (green) antibodies. PKC activation was assessed by phosphorylation of the downstream substrate, ERK using p44/42 Erk1/2 antibody. Arrows denote non-specific antibody background. **c** FAM126A phosphorylation at Ser485 was quantified as the ratio of total pFAM126A S485 signal/ total FLAG signal relative to DMSO treated FLAG-FAM126A WT. Data are the mean ± SEM (n=5). * *p* <0.05, ** *p*<0.01, *** *p*< 0.001, **** *p* <0.0001 using one-way ANOVAs with Dunnett’s multiple comparison tests.

Next, we used anti-pFAM126A S485 to probe FAM126A phosphorylation in cells under different signaling conditions. Although direct phosphorylation of the PI4KA complex has not been demonstrated, a recent study identified PKC as a possible regulator of this complex and showed that PMA activates PI4P production at the PM which is blocked by BIM, a PKC inhibitor ^28^. Therefore, we monitored FAM126A phosphorylation with anti-pFAM126A S485 under similar conditions by treating cells that co-expressed FLAG-FAM126A (WT or ASASAA mutant), TTC7B and EFRB upon treatment with combinations of a CN inhibitor (FK506), a PKC activator (PMA) and a PKC inhibitor (BIM). By examining the total intensity of anti-pFAM126 S485 signal (forms PI and PII), we made the following observations (Fig. 5c): first, for cells expressing FAM126A^WT^, addition of FK506 significantly increased Ser485 phosphorylation under all conditions (alone or together with PMA, PMA+BIM). Second, compared to FAM126A^WT^, cells expressing FAM126A^ASASAA^ showed higher levels of Ser485 phosphorylation under all conditions and inhibiting CN with FK506 had no further effect as expected for this CN-binding impaired mutant. Together, these findings show that S485 phosphorylation is CN-regulated. Furthermore, addition of PMA did not enhance S485 phosphorylation. Next, we focused on shifts in electrophoretic mobility of FAM126A (Fig. 5b). Interestingly, for both FAM126A proteins (WT and ASASAA mutant), treatment with PMA caused an electrophoretic shift from the dephosphorylated form (deP) to PI, likely due to phosphorylation of residues other than Ser485. Importantly, this PMA-induced shift was suppressed by BIM. In sum, these findings demonstrate PxIxIT-dependent regulation of FAM126A phosphorylation at Ser485 by CN *in vivo* and establish FAM126A as a novel CN substrate. These data also reveal PMA-induced phosphorylation of FAM126A, at a distinct site, likely by PM-localized PKC, which might be the molecular basis of the reported regulatory role for PKC in PM PI4P synthesis ^28^.

### CN regulates PI4P synthesis by the PI4KA complex

Having shown that CN interacts with the PI4KA complex and that FAM126A is a CN substrate, we next investigated whether CN regulates the assembly and/or activity of this complex. First, we examined interaction of the cytosolic heterotimer, PI4KA/TTCB/FAM126A, with the membrane anchor EFR3B in the presence of FAM126A^WT^ or CN-binding defective FAM126A^ASASAA^. Immunopurification of EFR3B-HA showed the same levels of co-purifying GFP-PI4KA, TTC7B-MYC or FLAG-FAM126A with FAM126A^WT^ or FAM126A^ASASAA^ (Fig. 6a, b). Furthermore, indirect immunofluorescence analyses of these cells verified that each component, especially GFP-PI4KA, localized to the PM with either FAM126A^WT^ or FAM126A^ASASAA^, indicating that the complex formed properly (Supplementary Fig. 5c). These findings suggest no CN-dependent regulation of complex formation via FAM126A.

**Figure 6:**
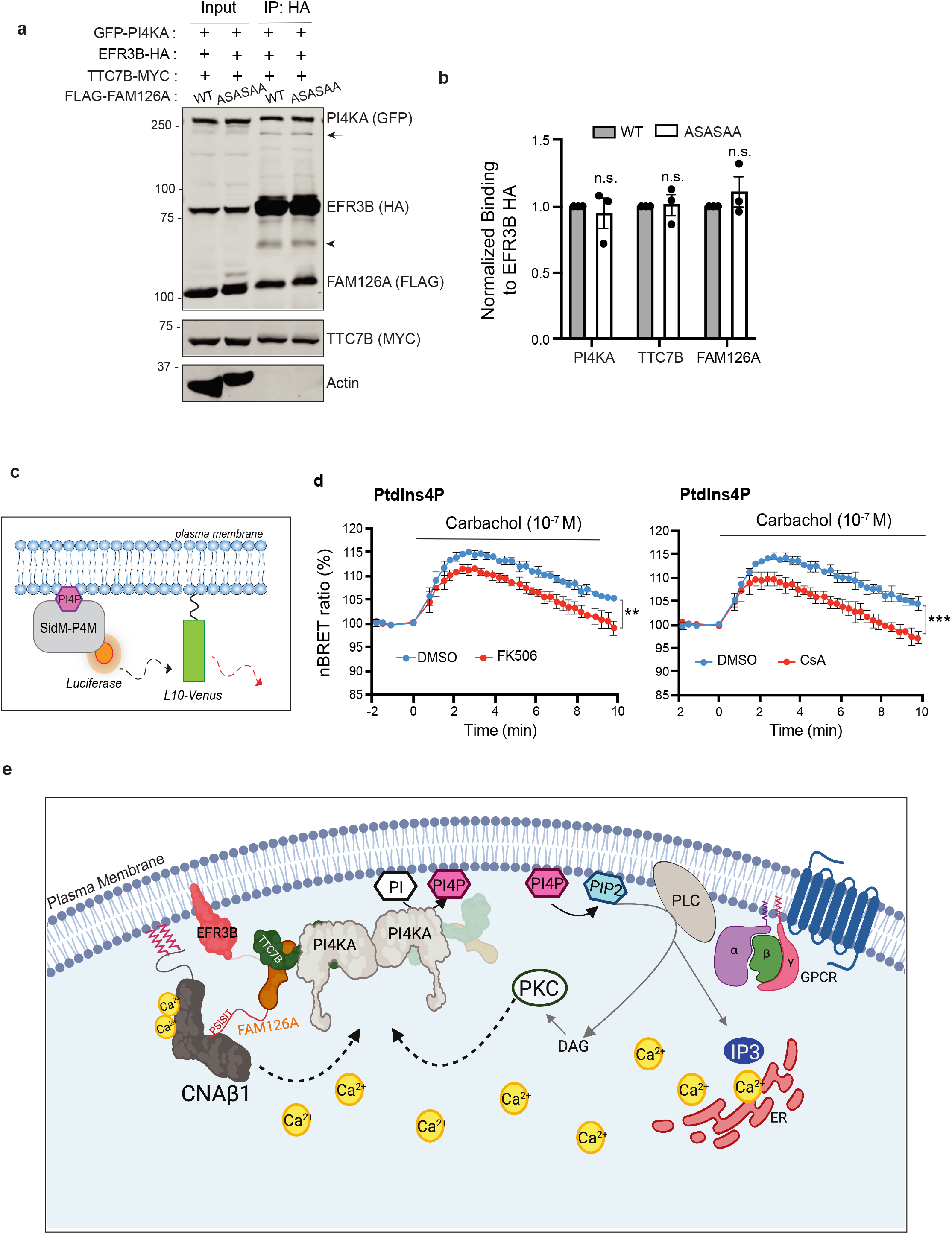
CN regulates PI4P synthesis by the PI4KA complex. **a**. Immunoblot analysis of the PI4KA complex components in anti-HA immunoprecipitates of HeLa cells expressing GFP-PI4KA, EFR3B-HA/TTC7B-MYC with WT or ASASAA mutant FLAG-FAM126A. Arrow points to the uncut P2A form. Arrowhead denotes non-specific antibody bands. **b** Co-purification of each component in with EFR3B-HA is quantified and normalized to input. Data are the mean ± SD (n=3). n.s. not significant. **c** Cartoon representation of the BRET pair used in the experiments. PI4P binding domain of the Legionella SidM protein (SidM-P4M) attached to Renilla Luciferase as the donor and the Venus targeted to the PM using the first 10 amino acids of Lck, L10, as the acceptor. **d** Normalized BRET ratios reflecting the changes in the PM PI4P levels in response of carbachol stimulation (10^-7^ M) in HEK293T cells transiently expressing muscarinic receptor, M3R, pre-treated with DMSO (blue), or CN inhibitors (red): FK506 (1 µM) (left) and Cyclosporin A, CsA (10µM) (right) for 1 h. Values are the mean ± SD (n=4). ** *p*= 0.0063, *** *p* =0.0004 **e** Model for CNAβ1 mediated regulation of the PI4KA complex that promotes the PM PI4P synthesis during GPCR signaling. The increased intracellular Ca^2+^ activates CN, and likely, PKC, which in turn regulate the PI4KA complex at the PM to promote PI4P and PI(4,5)P_2_ replenishment.

Next, we explored whether CN regulates PM PI4P synthesis carried out by the PI4KA complex using a previously established bioluminescence resonance energy transfer (BRET) assay that monitors PI4P levels at the PM in live cells during signaling ^28^. For this assay, the energy donor (luciferase) is fused to the PI4P binding domain, P4M, of the *Legionella* SidM protein ^57^, the energy acceptor (Venus) is attached to the PM-targeting sequence from Lck (first ten amino acids, L10) (Fig. 6c), and both proteins are expressed in HEK293 cells that express the Gq-coupled muscarinic receptor, M_3_R. As previously reported, PM PI4P levels transiently increased in control cells (blue lines, Fig. 6d) following addition of the M_3_R ligand, carbachol (10^-7^ M), due to activation of the PI4KA complex ^28^. Excitingly, pre-treatment of these cells with CN inhibitors, FK506 (1 µM) or CsA (10 µM), significantly reduced the level of PI4P produced (red lines, Fig. 6d) consistent with our hypothesis that CN regulates PI4KA complex activity under Ca^2+^ signaling conditions.

In summary, our findings lead us to propose the following model: Signaling from a Gq-coupled GPCR generates an intracellular Ca^2+^ signal that activates CN, and likely PKC, which in turn stimulate the PI4KA complex at the PM to promote the PI4P replenishment and thus generating the PI(4,5)P_2_ pools required for sustained signaling (Fig. 6e). Our work highlights CNAβ1 as a newly identified interaction partner of the PI4KA complex, shows that CN inhibitors alter PI4P production at the PM during signaling, and warrants further investigation into the phosphorylation state of complex components, especially FAM126A and PI4KA, through which CN might be regulating PI4KA activity.

## Discussion

In this study we aimed to discover CN signaling pathways that are regulated by the naturally occurring but understudied CN isoform, CNAβ1, which is conserved among vertebrates and broadly expressed ^13,14,15^. This isoform differs from canonical CNAβ2 only in its 40 C-terminal residues ^14^, which confer distinct enzymatic regulation to CNAβ1 through an LxVP-type autoinhibitory sequence (LAVP), that we previously characterized ^15^. Here we show that the CNAβ1 tail is dually palmitoylated, making CNAβ1 the only known form of CN that directly associates with the PM and Golgi. By contrast, canonical CN isoforms access only select PM proteins that either contain CN binding sites in their cytosolic domains (e.g. NHE1, TRESK) ^58, 59^ or associate with membrane-anchored scaffolds such as AKAP79 ^60^. This unique localization determines CNAβ1 substrate specificity including its interaction with all four members of the protein complex that synthesizes the critical phospholipid, PI4P at the PM. We demonstrate that FAM126A, the regulatory component of this complex, is phosphorylated at the PM, directly binds CN, and contains at least one CN-regulated phosphorylation site. These findings led us to discover a hitherto unknown role for CN in regulating PI4P synthesis at the PM during GPCR signaling. The CNAβ1 isoform is ideally positioned to carry out this regulation.

Our finding that CNAβ1 is dynamically palmitoylated has several interesting implications for its regulation *in vivo.* First, the ability of CNAβ1 to access membrane-associated substrates and hence carry out its functions may be controlled by the palmitoyl transferases (DHHCs) and depalmitoylases that act on it, as has been shown for other signaling enzymes including RAS and LCK ^61, 62^. Second, we speculate that the palmitoylation-driven binding of the autoinhibitory CNAβ1 tail to membranes may be necessary to fully activate this variant which is only partially activated by Ca^2+^ and CaM *in vitro* ^15^. Thus, examining the enzymes that modify CNAβ1 lipidation will be key for understanding how CNAβ1 is controlled physiologically. Here we show that a membrane-localized thioesterase ABHD17A, which regulates H/N-RAS ^44^, also catalyzes the depalmitoylation of CNAβ1 causing it to redistribute from the PM to the cytosol and the Golgi. However, mechanisms that control the activity of ABHD17A are yet to be identified. Furthermore, determining which of the 23 DHHCs encoded in human genome act on CNAβ1 may provide insights into where and when palmitoylation takes place, as these enzymes exhibit distinct patterns of localization and regulation ^63^. In sum, our findings lay the groundwork for further investigation into the role of dynamic palmitoylation in controlling CNAβ1 localization and/or enzymatic activity, which may also provide tools to specifically regulate its functions.

Our investigations are the first to identify CN as a regulator of the PI4KA complex composed of PI4KA, TTC7, FAM126A and EFR3, and highlight major gaps in our knowledge of how this important complex is regulated. Production and maintenance of PM PI4P levels are physiologically critical as evidenced by the wide range of diseases caused by mutations in complex components ranging from neurological (PI4KA), immune and gastrointestinal (TTC7) defects, to hypomyelination and congenital cataracts (FAM126A) ^30, 34^. Phosphorylation regulates assembly of the PI4KA complex in yeast; however, in mammals, little is known about how the assembly or the activity of this complex is modulated. Our interaction studies, including HDX-MS analysis, uncovered potential contacts between CN and multiple PI4KA complex members, and confirmed direct binding to a PxIxIT motif in the C-terminal tail of FAM126A. This tail is completely unstructured and shows no interaction with TTC7 or PI4KA ^30^, but inhibits PI4KA activity *in vitro* through an unknown mechanism ^35^. Our results provide the first insights into this regulation by demonstrating that CN binds to and modulates the phosphorylation of at least one site in the FAM126A tail (Ser485) in cells. Further studies are required to comprehensively map CN-regulated phosphosites in PI4KA complex members, identify relevant PM-associated kinases, and to assess the functional consequences of these modifications.

Lastly, our discovery that CN inhibitors reduce PI4P production at the PM induced during Ca^2+^ signaling from Gq-coupled GPCRs suggests that a positive feedback loop exists through which PKC and CN (presumably CNAβ1), regulate the phosphorylation of the PI4KA complex to stimulate its activity and ensure a continued supply of the precursor PI(4,5)P_2_ (Fig. 6e). Evidence for PKC involvement in this stimulation ^28^ is consistent with the CN-independent, PKC-regulated phosphorylation-shift we observed in FAM126A (Fig. 5b). Rigorously testing this model, however, is extremely challenging due to the complete lack of knowledge about how this large, minimally expressed complex that apparently undergoes extensive allosteric rearrangements ^35^, is regulated in cells. To date, our attempts to identify changes in PI4P levels or synthesis rates caused by mutations in the FAM126A PxIxIT site or Ser485 have been unsuccessful using overexpression of the proteins (data not shown). This could be due to limitations in the experimental set up (HEK293 cells overexpressing all PI4KA complex members), or because solely altering FAM126A may not be sufficient to perturb CN-dependent regulation of the complex. Regardless, our work breaks new ground by establishing that CNAβ1 preferentially interacts with the PI4KA complex at the PM and by suggesting FAM126A as the first substrate for CNAβ1.

Insights into the physiological functions of CNAβ1 come from studies that overexpress or more recently, delete the CNAβ1 isoform in mice. These knock-out mice are viable, but develop cardiac hypertrophy, possibly due to disruptions in mTORC2/AKT signaling and serine one-carbon metabolism ^24^. However, the precise molecular mechanisms underlying these pathologies and whether any of these phenotypes relate to PI4KA complex regulation remain to be determined. Notably, some reports indicate that mTORC2 activity toward AKT takes place at the PM and depends on the PH-domain containing targeting subunit, mSIN1, which binds to phosphoinositides ^64–66^. Furthermore, the interactors identified here suggest that CNAβ1 regulates multiple substrates throughout the body. Comprehensive identification of these targets as well as the regulatory mechanisms that control CNAβ1 activity *in vivo* promise to shed new light on Ca^2+^ and CN-regulated pathways and their possible perturbation in patients undergoing long term treatment with CN inhibitors, CsA or FK5006/Tacrolimus.

## Methods

### Sequence alignments

ClustalW was used to create all sequence alignments using Jalview. The following species are used in Fig. 1b: *Homo sapiens* (human, Q5F2F8), *Pan troglodytes* (chimpanzee, A0A2J8NUG2), *Sus scrofa* (pig, A0A480QFW6), *Desmodus rotundu*s (Bat, K9ISS2), *Mus musculus* (mouse, Q3UXV4), *Callorhinchus milii* (ghost shark, V9KGC1) and *Xenopus tropicalis* (western clawed frog, A0A6I8R6A9). The following species are used for Supplementary Fig. 4a: *Homo sapiens* (human, Q9BYI3), *Gorilla gorilla* (gorilla, A0A2I2YR80), *Macaca mulatta* (monkey, H9ZEG3), *Sus scrofa* (pig, I3LJX1), *Felis catus* (cat, M3WEC3), *Bos taurus* (bovine, E1BFZ6), *Mus musculus* (mouse, Q6P9N1), *Gallus gallus* (chicken, Q5ZM13*), Xenopus tropicalis* (western clawed frog, F7EHL4), *Callorhinchus milii* (ghost shark, A0A4W3JDV9), *Danio rerio* (zebrafish, Q6P121).

### Cell culture and transfection

HeLa, COS-7 and HEK 293T cells were grown at 37 °C in a 5% CO_2_ atmosphere in cell culture medium (Dulbecco’s modified Eaglès medium (CA 10-013, Sigma-Aldrich) supplemented with 10% fetal bovine serum (FBS, Benchmark^TM^ Gemini Bio Products). HEK 293 inducible cell lines were generated by transfection of Flp-In T-REx parental cells (obtained from the Gingras Lab) with flippase (pOG44) and indicated plasmids in the pCDNA5/FRT vector, followed by selection using 200 μg/ml hygromycin B (Sigma-Aldrich). HEK 293 Flip-In T-REx cell lines were maintained in cell culture medium supplemented with 200 μg/ml hygromycin B, 3 μg/ml blasticidin S Hydrochloride (RPI) and induced with 10ng/ml doxycycline (Sigma-Aldrich). HEK 293T and HeLa cells were gifts from the Skotheim lab. COS-7 cells were purchased from ATCC (CRL-1651). Cells were transfected as indicated in each experiment using jetOPTIMUS (VWR) as per the manufacturer’s instructions.

### Plasmids

DNAs encoding the human CNAβ1(1-496), CNAβ2(1-524) were subcloned into a pcDNA5 expression vector which encodes an N-terminal FLAG tag or GFP tag. FAM126A cDNA received from Addgene, was subcloned into pcDNA5 with N-terminal FLAG tag, between BamHI and XhoI sites. Variants of CNAβ1 (CNAβ1^C483S^, CNAβ1^C493S^ CNAβ1^C2S^) and FAM126A (FAM126A^ASASAA^, FAM126A^S485A^, FAM126A^ASASAA+S485A^) were generated using the Quickchange (Agilent) site-directed mutagenesis kit. Plasmids containing the DNA encoding ABHD17A (WT and S190A mutant), APT1, APT2 and Venus-RIT were gifts from the Conibear lab. CNA/CNB plasmid (residues 2-391 of human CNA alpha isoform and human CNB isoform 1) tandemly fused in pGEX6P3 (which encodes N -terminal GST tag) for protein purification and use in HDX-MS experiments were cloned as described before ^6^. 6x His-CNA (residues 1-391 of human CNA alpha isoform and human CNB isoform 1 tandemly fused in, p11 vector) used in in vitro peptide binding assays, was cloned as described before^21^ His-CN NIR (^330^NIR^332^ -AAA mutations) generated using site-directed mutagenesis using His-CN WT as template. Plasmids encoding human PI4KIIIα were gifts of the Balla lab. Plasmids encoding human EFR3B and TTC7B, both C-terminally tagged, were gifts of the De Camilli lab. EFR3BHA_T2A_TTCBMYC (or GFP)_P2A_FLAG FAM126A plasmid was generated in between HindIII and NotI site of pcDNA3.1 vector. The DNA sequence that encodes viral T2A (GSGEGRGSLLTCGDVEENPGP) was subcloned to the 5’ end of the EFR3B-HA sequence, TTC7B-MYC was then subcloned in frame to the T2A sequence. FLAG-FAM126A with DNA sequence encoding for P2A (GSGATNFSLLKQAGDVEENPGP) was cloned in frame, at the 5’ end of the TTC7B-MYC sequence.

### Antibodies

Antibodies used in each experiment are listed for each methods section with their working dilutions listed in parentheses. The phosphospecific antibody against Serine 485 site in FAM126A was manufactured by 21st Century Biochemicals as follows: A peptide corresponding to the sequence Hydrazine-Ahx-ANRFSAC[pS]LQEEKLI-amide was manufactured by Fmoc chemistry, HPLC purified to >90%, and its mass and sequence were verified by nanospray MS and CID MS/MS, respectively. The peptides, along with carrier proteins and adjuvant, were injected into New Zealand White rabbits using an initial CFA injection, followed by IFA injections. A production bleed was then taken from each of the rabbits. Sera were passed multiple times over a hydrazine reactive resin which was linked to the immunogen peptides, then rinsed with both salt and phosphate buffers. The antibody fractions were collected using an acidic elution buffer and immediately neutralized before a two-stage dialysis into PBS buffer, pH 7.2. The antibody concentration was determined using a spectrophotometer (A280). The purified antibodies were then passed multiple times over a hydrazine reactive resin, which is linked with the unmodified peptides (those not injected). These immunodepletion steps were done to remove any non-specific/phospho-independent antibodies. The final antibodies were then buffered in a PBS/50% glycerol buffer, pH 7.2 and the final concentration was calculated using a spectrophotometer (A280).

### Immunofluorescence, microscopy and image analysis

HeLa or COS-7 cells were grown on 12 mm, #1.5H glass coverslips (ThorLabs). 24 h post-transfection, cells were washed with 1X PBS and fixed in 4% paraformaldehyde (PFA) solution (diluted from 16% PFA, Electron Microscopy Sciences) in PBS for 15 min. Cells were washed thrice with PBS and permeabilized for 5 min in block buffer (1x PBS with 0.2 M Glycine, 2.5% FBS) with 0.1% Triton X-100. Cells were then incubated in block buffer without detergent for 30 min. Coverslips were incubated with primary antibodies diluted in block buffer (without detergent) for 1 h, washed multiple times with 1x PBS followed by incubation with secondary antibodies for 1 h at room temperature. Coverslips were washed again and mounted using Prolong Diamond Antifade mountant (Thermo Fisher). Images were acquired on a single z-plane on Lionheart^TM^ FX automated widefield microscope with a 20X Plan Fluorite WD 6.6 NP 0.45 objective. For Fig. 1e and Supplementary Fig. 1e, primary antibodies used: mouse anti-FLAG, M2 (1:500, Sigma-Aldrich, F1804) and rabbit anti-GM130, D6B1 (1:400, Cell Signaling Technologies, 12480). Secondary antibodies used: Anti-mouse Alexa Fluor 647 (1:500, Invitrogen) and anti-rabbit Brilliant Violet 421 (1:100, Biolegend). YFP (500 nm), Texas Red (590 nm) and DAPI (350 nm) filter cubes were used to image Venus, FLAG and GM130 respectively. For Fig. 2f: GFP (465 nm), Texas Red and DAPI filter cubes were used to image GFP, FLAG and GM130, respectively.

#### Image analysis

Image analyses were performed in ImageJ, FIJI. The EzColocalization plugin for FIJI was used for co-localization analysis in Fig. 1f and Supplementary Fig. 1f to determine the Pearson correlation coefficients ^67^. For PM localization, a binary mask was generated from the thresholded Venus-RIT channel and saved as a selection (outer) to measure total signal intensity cell. The second mask was produced by five iterations of erosion function and subtracted from the outer mask using image calculator. The resulting mask (Supplementary Fig. 1d) was converted to a selection and used to measure the PM signal intensity.

### Detergent-assisted subcellular fractionation

COS-7 cells were seeded onto 60 mm plates and transfected with FLAG-CNAβ2, FLAG-CNAβ1 (WT or C483S, C493S, C483/C493S) or EFR3B-FLAG at 80% confluency. 48 h post transfection, cells were rinsed, harvested and pellets were snap-frozen in liquid nitrogen. Pellets were resuspended in 200 μl digitonin buffer (10 mM HEPES pH 6.8, 100 mM NaCl, 300 mM sucrose, 3 mM MgCl_2_, 5 mM EDTA and 0.015% Digitonin) supplemented with protease inhibitors by pipetting up and down followed by rotating at 4°C for 15 min. Input (6%) was taken as input prior to centrifugation at 2,000g for 20 min. The supernatant was carefully removed and spun at 16,000g for 5 min to remove any contamination from the pellet fraction. The supernatant was saved as the cytosol fraction. The pellet was washed 2x with ice-cold PBS and resuspended in 200 μl Triton X-100 buffer (HEPES pH 7.5, 100 mM NaCl, 300 mM sucrose, 3 mM MgCl_2_, 3 mM EDTA, 1% Triton X-100) supplemented with protease inhibitors. Pellets were lysed for 30min by rotating at 4°C followed centrifugation at 7,000g for 10 min. Supernatant saved as the membrane fraction. The clarified supernatant is saved as the cytosol fraction. Inputs and equal volumes (6%) of the cytosol and membrane fractions were mixed with 6X SDS sample buffer, heated to 95°C for 5 min and resolved by SDS-PAGE followed by western blotting. Primary antibodies used: anti-FLAG (1:2,500; Sigma F3165), rabbit anti-calnexin (1:3,000 ADI-SPA-865, Enzo Life Sciences) and anti-Gapdh (1:20,000, 1E6D9, Proteintech). After incubation with secondary Li-Cor antibodies, blots were imaged with the Li-Cor Odyssey imaging system. Enrichment in cytosol fraction was quantified as FLAG signal /Gapdh signal in cytosol fraction normalized to FLAG signal/Gapdh signal in Inputs. Similarly for membrane enrichment, FLAG signal /Calnexin signal in membrane fraction normalized to FLAG signal / Calnexin signal in Input. Statistical analysis was performed using GraphPad.

### Acyl-Resin Assisted Capture (Acyl-RAC)

The acyl-RAC protocol was performed as described previously^40^ with minor changes. In brief, COS-7 cells were seeded on 60mm plates and transfected at 70% confluency with FLAG-CNAβ2, FLAG-CNAβ1 (WT or C483S, C493S, C483/C493S) or EFR3B-FLAG using JetOptimus. 48 hours following transfection, cells were harvested in ice-cold PBS and snap frozen in liquid nitrogen. Pellets were lysed in TAE lysis buffer (50 mM TEA pH 7.3, 150 mM NaCl, 2.5% SDS) supplemented with 1 mM PMSF and protease inhibitors, vortexed briefly and incubated at 37°C for 20 min with constant gentle agitation. Lysates were subjected to fine needle aspiration with sterile 27.5-gauge needle and clarified by centrifugation (16,000g for 20 min). 400 μg of each lysate was diluted to 2 mg/ml with lysis buffer and incubated with 10 mM TCEP (646547, Sigma-Aldrich) for 30 min, nutating at room temperature. 25 mM NEM (N-ethylmaleimide, 40526, Alfa Aesar) was then added to the mix and incubated by gentle mixing at 40°C for 2 h to block free thiols. NEM was removed by acetone precipitation by adding four volumes of ice-cold acetone. Proteins were allowed to precipitate at -20°C overnight. Following centrifugation of the solution at 16,000g for 15 min, the pellets were extensively washed with 70% acetone and the pellets were airdried for 5 min at room temperature. Pellets were resuspended in 200 μl of binding buffer (50 mM TEA pH 7.3, 150 mM NaCl, 1 mM EDTA, 1% SDS. 0.2% Triton X-100) by heating at 40°C with frequent mixing. Approximately 20 μl from each sample was taken as input and the rest were split into two 1.5-ml microcentrifuge tubes. To capture S-palmitoylated proteins, 40 μl prewashed thiopropyl Sepharose 6b (T8387, Sigma-Aldrich, prepared fresh) was added to samples in the presence of either 0.75 M NH_2_OH (from 2.5 M stock, pH 7.5, freshly diluted from Hydroxylamine solution (467804, Sigma-Aldrich)) or binding buffer (without SDS and EDTA for the negative control). Binding reactions were carried out on a rotator at 30°C for 4 h. Resins were washed 4-5x with binding buffer, 5 min each, and proteins were eluted in 30 μl binding buffer supplemented with 50 mM DTT shaking at 30°C for 30 min. 6x SDS sample buffer was added to the samples followed by heating to 95°C for 5 min. Inputs and eluates were separated by SDS-PAGE and transferred to nitrocellulose for Western blotting with mouse anti-FLAG (1:2,500; Sigma F3165) and rabbit anti-calnexin (1:3,000 ADI-SPA-865, Enzo Life Sciences) antibodies. After incubation with secondary Li-Cor antibodies, blots were imaged with the Li-Cor Odyssey imaging system.

### Acyl-PEG Exchange (APE)

COS-7 cells were transfected, lysed, and subjected to reductive alkylation with TCEP and NEM as described in Acyl-RAC protocol. Following alkylation of total lysate (300-400 μg) proteins were precipitated with four volumes of ice-cold Acetone at -20°C overnight. The pellets were washed extensively with 70% Acetone and air dried for 5min. The pellets were resuspended in 72 μl TEA buffer pH 7.3, with 4% SDS (50 mM TEA, 150 mM NaCl, 0.2% Triton X-100, 4 mM EDTA) by heating to 40°C for an hour with constant mixing. Lysate was clarified by centrifugation at 16,000g for 5 min. Approximately 7 μl (10%) from each sample was removed as input, the rest was split into two 30 μl aliquots. For NH_2_OH treated sample, 36μl NH_2_OH (2.5 M stock) was added and brought up to 120 μl with TEA buffer with 0.2% Triton X-100 (50 mM TEA, 150 mM NaCl). For negative control not treated with NH_2_OH, 90 μl TEA buffer with 0.2% Triton X-100 was added. After incubation at 30°C for 1 h on a rotator, proteins were precipitated using methanol-chloroform-H_2_O, briefly air dried and resuspended in 30 μl TEA buffer with 4% SDS, 50 mM TEA, 150 mM NACl, 0.2% Triton X-100, 4 mM EDTA by gentle mixing at 40°C. Each sample was treated with 90 μl TEA buffer with 1.33 mM mPEG-Mal (Methoxypolyethylene glycol maleimide, 5 kDa, 63187 Sigma-Aldrich) for a final concentration of 1 mM mPEG-Mal. Samples were incubated for 2h at RT with nutation before a final methanol-chloroform-H_2_O precipitation. The pellets were resuspended in 50 μl TAE lysis buffer (50 mM TEA pH 7.3, 150 mM NaCl, 2.5% SDS) and 10 μl 6X SDS sample buffer was added before heating the sampled for 5min at 95°C. Typically 14μl of each sample was separated by SDS-PAGE and analyzed by western blot with FLAG and Calnexin antibodies. After incubation with secondary Li-Cor antibodies, blots were imaged with the Li-Cor Odyssey imaging system.

### Pulse-chase metabolic labeling with Palmostatin B

COS-7 cells were transfected with cDNA encoding FLAG-CNAβ1 using Lipofectamine 2000 as per manufacturer’s instructions. Twenty hours following transfection, cells were washed in phosphate-buffered saline (PBS) and starved in methionine-free DMEM containing 5% charcoal-filtered FBS (Life Technologies), supplemented with 1 mM L-glutamine and 1 mM sodium pyruvate for 1 h. Cells were then briefly washed in PBS then labeled with 30 μM 17-ODYA and 50 μM L-AHA for 2 h in this media. Labelling media was removed, cells were washed twice in PBS before chasing in complete DMEM supplemented with 10% FBS and 300 μM palmitic acid. Palmostatin B (Palm B) or DMSO (vehicle) were added at chase time 0 and Palm B was replaced every hour. At indicated time points, cells were washed twice in PBS and frozen at -80°C until processing. Cells were lysed with 500 μl triethanolamine (TEA) lysis buffer (1% Triton X-100, 150 mM NaCl, 50 mM TEA pH 7.4, 100xEDTA-free Halt Protease Inhibitor [Life Technologies]). The lysates were transferred to 1.5-ml Eppendorf tubes (Corning), vigorously shaken while placed on ice in between each agitation. Lysates were cleared by centrifugation at 13,000 g for 15 min at 4°C. Solubilized proteins in the supernatant were quantified using Bicinchoninic acid (BCA) assay (Life Technologies), 650μg-1mg of the lysate was added to Protein A-Sepharose beads (GE Healthcare) pre-incubated for 3-7 h with rabbit anti-FLAG antibody (Sigma-Aldrich) at 4°C. Immunoprecipitations were carried out overnight rotating at 4°C.

#### Sequential on-bead CuAAC/click chemistry

Sequential on-bead click chemistry of immunoprecipitated 17-ODYA/L-AHA-labeled proteins was carried out as previously described ^44^ with minor modifications. After immunoprecipitation, Sepharose beads were washed thrice in RIPA buffer, and on-bead conjugation of AF647 to 17-ODYA was carried out for 1 h at room temperature in 50 μl of freshly mixed click chemistry reaction mixture containing 1 mM TCEP, 1 mM CuSO_4_.5H_2_O, 100 μM TBTA, and 100 mM AF647-azide in PBS. After three washes in 500 μl ice-cold RIPA buffer, conjugation of AF488 to L-AHA was carried out for 1 h at room temperature in 50 μl click-chemistry reaction mixture containing 1 mM TCEP, 1 mM CuSO4.5H2O, 100 μM TBTA, and 100 mM AF488-alkyne in RIPA buffer. Beads were washed thrice with RIPA buffer and resuspended in 10 μl SDS buffer (150 mM NaCl, 4% SDS, 50 mM TEA pH7.4), 4.35 μl 4X SDS-sample buffer (8% SDS, 4% Bromophenol Blue, 200 mM Tris-HCl pH 6.8, 40% Glycerol), and 0.65 μl 2-mercaptoethanol. Samples were heated for 5 min at 90°C and separated on 10% tris-glycine SDS-PAGE gels for subsequent in-gel fluorescence analyses. A Typhoon Trio scanner (GE Healthcare) was used to measure in-gel fluorescence of SDS–PAGE gels: AF488 signals were acquired using the blue laser (excitation 488 nm) with a 520BP40 emission filter, AF647 signals were acquired using the red laser (excitation 633 nm) with a 670BP30 emission filter. Signals were acquired in the linear range and quantified using the ImageQuant TL7.0 software (GE Healthcare). For pulse-chase analyses, the ratio of palmitoylated substrates:total newly synthesized substrates were calculated as AF647/AF488 values at each time point, normalized to the value at T=0.

### Determination of CNAβ1 palmitoylation in the presence of thioesterases

COS-7 cells were seeded onto 60mm plates and transfected with GFP-CNAβ1 together with vector, ABHD17A-FLAG (WT or S190A mutant), FLAG-APT2 or mCherry-APT1. 24 h post-transfection, media was replaced with DMEM containing 2% FBS and labelled with 30 μM 17-ODYA (17-Octadecynoic Acid,34450, Cayman Chemicals) or DMSO for 3hr at 37°C incubator. Cells for rinsed thrice with ice-cold PBS, harvested and pellets were snap-frozen in liquid nitrogen. Pellets were then lysed in TEA lysis buffer ( 50 mM TEA pH 7.4, 150 mM NaCl, 1% Triton X-100, 1 mM PMSF) supplemented with protease inhibitors by rotating for 20min at 4°C. Lysates were subjected to fine-needle aspiration using sterile 27G syringe and clarified by spinning down at 16,000g for 15 min. 300-400 μg of each lysate was adjusted to 1mg/ml with TAE lysis buffer and bound to 10μl pre-washed GFP-trap magnetic particles in for 1-2h rotating end-over-end at 4°C. Input (5%) was taken prior to bead binding. Beads were washed thrice in modified RIPA buffer (50 mM TAE pH 7.4, 150 mM NaCl, 1% Triton X-100, 1% sodium deoxycholate, 0.1% SDS). Proteins bound to beads were conjugated to azide-biotin in 50μl PBS with click chemistry reactants for 1 h at RT with constant agitation. Click chemistry reactants were freshly prepared as a 5X master mix that consists of 0.5 M biotin-azide (Biotin-Picolyl azide, 1167, Click Chemistry Tools), 5 mM TCEP, 0.5 mM TBTA (Tris[(1-benzyl-1H-1,2,3-Triazol-4-yl)methyl]amine, Sigma-Aldrich) and 5 mM CuSO_4_.5H_2_O. Beads were washed thrice in modified RIPA buffer and proteins were eluted by boiling in 2X SDS sample buffer before resolving with SDS-PAGE. Anti-GFP (1:4,000, Living Colors, 632380, Clontech) was used to probe for GFP-CNAβ1, biotin incorporation was detected using fluorophore conjugated Streptavidin antibody (Licor IRDye 800CW Steptavidin, LI-COR Biosciences). Amount of ABHD17A and APT2 was probed using FLAG (1:2,500; F3165, Sigma-Aldrich) antibody and APT1 was detected using for anti-RFP (1:3,000; 22904, Rockland Inc.) Level of GFP-CNAβ1 palmitoylation was quantified as streptavidin signal normalized to bound GFP signal. Statistical analyses were performed in GraphPad.

### Affinity purification coupled to Mass Spectrometry (AP/MS) Analyses

#### Stable Cell Line Generation

Human [taxid:9606] cells [Flp-In T-REx 293 cells], were transfected in a 6-well format with 0.2 μg of tagged DNA [pcDNA5-FLAG-protein] and 2 μg pOG44 (OpenFreezer V4134), using lipofectamine PLUS (Invitrogen), according to the manufacturer’s instructions. On day 2, cells were trypsinized, and seeded into 10 cm plates. On day 3, the medium is replaced with DMEM containing 5% fetal bovine serum, 5% calf serum, 100 units/ml penicillin/streptomycin, and 200 ug/ml hygromycin. Medium was replaced every 2-4 days until non-transfected cells died and isolated clones were ∼1-2 mm in diameter (13-15 days). Pools of cells were generated by trypsinization of the entire plate and replating in fresh selection medium (the size of the plate was dictated by the number and size of initial colonies). Pools were amplified to one 15cm plate. From this plate, cells were trypsinized (volume = 8 ml) and replated in five 15cm plates. A frozen stock was generated from the plate when cells reached ∼80% confluence. Cells at ∼60-70% confluence were induced with 1 μg/ml tetracycline for 24 hours. Subconfluent cells (∼85-95% confluent) were harvested as follows: medium was drained from the plate, 0.5 ml ice-cold PBS was added, and the cells were scraped (using a silicon cake spatula) and transferred to a 1.5 ml Eppendorf tube on ice. Cells were collected by centrifugation (5 min, 1500 g, 4°C), the PBS aspirated, and cells resuspended in 1 ml ice-cold PBS prior to centrifugation (5 min, 1,500 g, 4°C). This step was repeated once more, the remaining PBS was aspirated, and the weight of the cell pellet was determined. Cell pellets were frozen on dry ice and transferred to -80°C until processing.

#### Affinity Purification

Cells were lysed by passive lysis assisted by freeze-thaw. Briefly, to the frozen cell pellet, a 1:4 pellet weight:volume ratio of ice-cold lysis buffer was added, and the frozen pellet was resuspended by pipetting up and down. The lysis buffer was 50 mM HEPES-NaOH pH 8.0, 100 mM KCl, 2 mM EDTA, 0.1% NP40, 10% glycerol, 1 mM PMSF, 1 mM DTT and Sigma protease inhibitor cocktail, P8340, 1:500. Tubes were frozen and thawed once by placing on dry ice for 5-10min, then incubated in a 37°C water bath with agitation until only a small amount of ice remained. Thawed samples were then put on ice, and the lysate transferred to 2 ml Eppendorf tubes. An aliquot (20 μl) was taken to monitor solubility. This aliquot was spun down, the supernatant transferred to a fresh tube, and 6 µl 4X Laemmli sample buffer added. The pellet was resuspendended in 26 µl 2X Laemmli sample buffer). The 2 ml tubes were centrifuged at 14,000 rpm for 20 min at 4°C, and the supernatant transferred to fresh 15 ml conical tubes. During centrifugation, anti-FLAG M2 magnetic beads (SIGMA) were prepared: 25 μl 50% slurry was aliquoted for each IP (two 150 mm plates), and the beads were washed in batch mode with 3 x 1 ml of lysis buffer. To the rest of the lysate, the equivalent of 12.5 μl packed FLAG M2 magnetic beads was added, and the mixture incubated 2 hours at 4°C with gentle agitation (nutator). Beads were pelleted by centrifugation (1,000 rpm for 1 min) and a 15 μl aliquot of the lysate post-IP was taken for analysis. Most of the supernatant was removed with a pipette, and the beads were transferred with ∼200 μl of lysis buffer to a fresh 1.7ml Eppendorf tube, magnetized for ∼30 s, and the remaining buffer was aspirated. Two washes with 1 ml lysis buffer and two washes with 20 mM Tris-HCl (pH 8.0) 2 mM CaCl_2_ were performed. Briefly, for each of these quick washes, the sample was demagnetized, resuspended by pipetting up and down in the wash buffer, remagnetized for ∼30 s, and the supernatant aspirated (a complete wash cycle takes between 1-2 min). After the last wash, most of the liquid was removed, the tube was spun briefly (1,000 rpm for 1 min), and the remaining drops were removed with a fine pipet.

#### Tryptic Digestion

The beads were resuspended in 5 μl of 20 mM Tris-HCl (pH 8.0). 500 ng of trypsin (Sigma Trypsin Singles, T7575; resuspended at 200ng/ul in Tris buffer) was added, and the mixture was incubated at 37°C with agitation for 4 hours. After this first incubation, the sample was magnetized and the supernatant transferred to a fresh tube. Another 500ng of trypsin was added, and the resulting sample was incubated at 37°C overnight (no agitation required). The next morning, formic acid was added to the sample to a final concentration of 2% (from a 50% stock solution).

#### Mass Spectrometry

Half the sample was used per analysis. A spray tip was formed on fused silica capillary column (0.75 μm ID, 350 μm OD) using a laser puller (program = 4; heat = 280, FIL = 0, VEL = 18, DEL = 200). 10 cm (±1 cm) of C18 reversed-phase material (Reprosil-Pur 120 C18-AQ, 3 μm) was packed in the column by pressure bomb (in MeOH). The column was then pre-equilibrated in buffer A (6 μl) before being connected in-line to a NanoLC-Ultra 2D plus HPLC system (Eksigent, Dublin, USA) coupled to an LTQ-Orbitrap Velos (Thermo Electron, Bremen, Germany) equipped with a nanoelectrospray ion source (Proxeon Biosystems, Odense, Denmark). The LTQ-Orbitrap Velos instrument under Xcalibur 2.0 was operated in the data dependent mode to automatically switch between MS and up to 10 subsequent MS/MS acquisitions. Buffer A was 100% H_2_O, 0.1% formic acid; buffer B was 100 ACN, 0.1% formic acid. The HPLC gradient program delivered the acetonitrile gradient over 125 min. For the first 20 minutes, the flow rate was of 400 μl/min at 2% B. The flow rate was then reduced to 200 μl/min and the fraction of solvent B increased in a linear fashion to 35% until min 95.5. Solvent B was then increased to 80% over 5 minutes and maintained at that level until 107 min. The mobile phase was then reduced to 2% B until the end of the run (125 min). The parameters for data dependent acquisition on the mass spectrometer were: 1 centroid MS (mass range 400-2000) followed by MS/MS on the 10 most abundant ions. General parameters were: activation type = CID, isolation width = 1 m/z, normalized collision energy = 35, activation Q = 0.25, activation time = 10 ms. For data dependent acquisition, minimum threshold was 500, the repeat count = 1, repeat duration = 30 s, exclusion size list = 500, exclusion duration = 30 s, exclusion mass width (by mass) = low 0.03, high 0.03.

#### Mass spectrometry data extraction

RAW mass spectrometry files were converted to mzXML using ProteoWizard (3.0.4468) and analyzed using the iProphet pipeline ^68^ implemented within ProHits ^69^ as follows. The database consisted of the human and adenovirus complements of the RefSeq protein database (version 57) supplemented with “common contaminants” from the Max Planck Institute (http://lotus1.gwdg.de/mpg/mmbc/maxquant_input.nsf/7994124a4298328fc125748d0048fee2/$FILE/contaminants.fasta) and the Global Proteome Machine (GPM; http://www.thegpm.org/crap/index.html). The search database consisted of forward and reversed sequences (labeled “DECOY”); in total 72,226 entries were searched. The search engines used were Mascot (2.3.02; Matrix Science) and Comet ^70^ (2012.01 rev.3) with trypsin specificity (two missed cleavages were allowed) and deamidation (NQ) and oxidation (M) as variable modifications. Charges of +2, +3 and +4 were allowed, and the parent mass tolerance was set at 12 ppm while the fragment bin tolerance was set at 0.6 amu. The resulting Comet and Mascot search results were individually processed by PeptideProphet ^71^ and peptides were assembled into proteins using parsimony rules first described in ProteinProphet ^72^ into a final iProphet protein output using the Trans-Proteomic Pipeline (TPP; Linux version, v0.0 Development trunk rev 0, Build 201303061711). TPP options were as follows: general options were -p0.05 -x20 -PPM -d“DECOY”, iProphet options were – ipPRIME and PeptideProphet options were –OpdP. All proteins with a minimal iProphet protein probability of 0.05 were parsed to the relational module of ProHits. Note that for analysis with SAINT, only proteins with iProphet protein probability ≥ 0.95 are considered. This corresponds to an estimated protein-level FDR of ∼0.5%. Statistical analysis was performed with SAINTexpress (with default parameters), using 38 biological replicates of FLAG-GFP (all from asynchronous HEK293 T-REx cells, all run on the Orbitrap Velos) as negative controls, including two samples run in tandem with the two biological replicates. Note that the negative control experiments were previously published ^73, 74^. High-confidence interaction partners can be found in Supplementary Table 1. All mass spectrometry data was deposited to ProteomeXchange through partner MassIVE and assigned the identifiers PXD026809 and MSV000087664, respectively. The dataset can be accessed prior to publication at ftp://MSV000087664@massive.ucsd.edu (password: calcineurin).

### Immunoprecipitations

For Fig. 3d, HEK 293 Flp-In T-REx cells expressing GFP vector alone, GFP-CNAβ2, GFP-CNAβ1 WT or C2S (C483/C493S) constructs were co-transfected with EFR3B HA_T2A_TTC7 MYC_P2A_FLAG FAM126A (WT) with GFP-PI4KA. 4 h post-transfection, the media were replaced with fresh media containing 10 ng/ml doxycycline and cells were induced for 24 h. Cells were rinsed with ice-cold PBS and harvested using a scraper. Harvested cells were pelleted by rotating at 3,500 rpm for 5 min and pellets were snap-frozen in liquid nitrogen and stored at -80°C. until use. Cell pellets were lysed in lysis buffer (50mM Tris, pH 7.5, 150 mM NaCl, 1% Triton X-100) supplemented with a protease and phosphatase inhibitor cocktail (Halt^TM^, ThermoFisher) and 250 U/ml benzonase for 30 min rotating end-over-end at 4°C and subjected to fine needle aspiration with sterile 27.5-gauge needle. Cell lysates were clarified by centrifugation at 16,000g for 20min and subjected to BCA assay to determine their protein concentrations. 600-800 µg of each lysate were adjusted to 1 mg/ml with binding buffer (50 mM Tris, pH 7.5, 150 mM NaCl, 0.5% Triton X-100) and bound to 40 μl Pierce anti-HA magnetic beads (ThermoFisher) for 4 h rotating at 4°C. Beads were washed thrice in binding buffer and co-precipitated proteins were eluted by boiling in 2X SDS sample buffer for 5min. Input (2%) and bound (100%) fractions were resolved by SDS-PAGE and immunoblotted with HA (1:2,000, H3663, Sigma-Aldrich), GFP (1:4,000, Living Colors, 632380, Clontech), MYC (1:3,000, 9B11, Cell Signaling Technologies) and β-Actin (1:3,000; 926-42210, Li-Cor Biosciences) antibodies followed by secondary Li-Cor antibodies. Blots were imaged with the Li-Cor Odyssey imaging system.

Binding of GFP-proteins was quantified as GFP signal/MYC bound signal normalized to the GFP signal/Actin Input signal. Statistical significance was determined using GraphPad. For Fig. 6a, HeLa cells co-transfected with EFR3B HA_T2A_TTC7 MYC_P2A_FLAG FAM126A (WT or ASASAA) and GFP-PI4KA constructs. 48h post transfection, cells were harvested, processed, and bound to anti-HA beads as described above. For Supplementary Fig. 3b, HEK293 Flp-In T-REx expressing GFP vector alone, GFP-CNAα, GFP-CNAβ2, GFP-CNAβ1 WT or C2S were transfected with FLAG-NFATC1. Cells were induced, harvested, and lysed as described above. 1000 μg of each lysate was bound to 15 μl pre-washed GFP-Trap magnetic beads (Bulldog Bio. Inc.) in 1 ml binding buffer for 2h, rotating end-over-end at 4°C. Beads were washed thrice in binding buffer and eluted by boiling in 2x SDS sample buffer for 5 min. Inputs (2%) and eluates were resolved by SDS-PAGE followed by western blotting. Binding was quantified as described above.

### Proximity-dependent biotin identification (BioID) analysis

HeLa cells were seeded onto 10cm plates and transfected at 80% confluence with MYC-BirA-CNAβ1. 24 hours post transfection, cells were passaged onto two 10 cm plates and let to grow overnight. The next day, cells were co-transfected with EFR3B HA_T2A_TTC7B GFP_P2A_ FLAG FAM126A (WT or ASASAA mutant) and HA-PI4KA. Four hours post-transfection, the media were replaced with fresh media containing 50 μM D-biotin (Sigma B-4501). After 16 hours of labeling, cells were collected and snap frozen in liquid nitrogen. Cells were lysed in RIPA buffer (150 mM NaCl, 1% Triton X-100, 0.5% Deoxycholate, 0.1% SDS, 50 mM Tris pH 8.0) supplemented with a protease and phosphatase inhibitor cocktail (Halt^TM^, ThermoFisher) and 250 U/ml benzonase (EMD Millipore) for 30 min rotating end-over-end at 4°C and subjected to fine needle aspiration with a sterile 27.5-gauge needle. Cell lysates were clarified by centrifugation (16,000g for 20 min). Solubilized protein concentration in the supernatant were quantified using BCA analysis. For each binding reaction, 1 mg of clarified lysate was incubated with 30 μl of pre-rinsed streptavidin magnetic particles (11641786001, Sigma-Aldrich) in 1 ml RIPA buffer for 16h, rotating at 4°C. An input aliquot (20 μl) was removed prior to bead addition. Beads were washed three times with 1ml RIPA buffer, rotating for 5 min each, and eluted in 2X sample buffer (10%SDS, 0.06% Bromophenol blue, 50% glycerol, 0.6 M DTT, 375 mM Tris-HCl pH 6.8). Inputs and bound (100%) samples were boiled and resolved by SDS-PAGE followed by western blotting with mouse FLAG (1:2,500; F3165, Sigma-Aldrich), rabbit MYC (1:2,000; 71D10, Cell Signaling), mouse HA (1:2,000, H3663, Sigma-Aldrich), mouse GFP (1:4,000, Living Colors, 632380, Clontech) and rabbit β-Actin (1:3,000; 926-42210, Li-Cor Biosciences) antibodies. Blots were imaged with Li-Cor Odyssey imaging system following incubation with secondary Li-Cor antibodies. Binding for each protein was quantified as their respective signals/MYC bound signal normalized to respective signals/Actin Input signal. Biotinylation of each protein in complex with WT FAM126A, is set to 1. Statistical significance was determined using GraphPad.

### In vitro peptide binding assays

#### Purification of Calcineurin

6xHis-tagged human calcineurin A (α isoform, truncated at residue 392), WT or ^330^NIR^332^-AAA mutant were expressed in tandem with the calcineurin B subunit in *E. coli* BL21 (DE3) cells (Invitrogen, USA) and cultured in LB medium containing carbenicillin (50 μg/ml) at 37°C to mid-log phase. Expression was induced with 1 mM IPTG at 16°C for 18 h. Cells were pelleted, washed and frozen at -80°C for at least 12 h. Thawed cell pellets were re-suspended in lysis buffer (50 mM Tris-HCl pH 7.5, 150 mM NaCl, 0.1% Tween 20, 1mM β-mercaptoethanol, protease inhibitors) and lysed by sonication using four, 1-minute pulses at 40% output. Extracts were clarified using two rounds of centrifugation (20,000 X g, 20 min) and then bound to 1 ml of Ni-NTA agarose beads (Invitrogen) in lysis buffer containing 5mM imidazole for 2-4 hr. at 4°C, in batch. Bound beads were loaded onto a column and washed with lysis buffer containing 20 mM imidazole and eluted with lysis buffer containing 300 mM imidazole, pH 7.5. Purified calcineurin heterodimer were dialyzed in buffer (50 mM Tris-HCl pH 7.5, 150 mM NaCl, 1 mM β-mercaptoethanol) and stored in 10% glycerol at -80°C.

#### Peptide purification

16mer peptides were fused to GST in vector pGEX-4T-3 and expressed in *E. coli* BL21 (DE3) (Invitrogen). Cells were grown at 37°C to mid-log phase and induced with 1 mM IPTG for 2 hr. Cell lysates were prepared using the EasyLyse^TM^ bacterial protein extract solution (Lucigen Corp. USA) or the CelLytic B reagent (Sigma, USA) according to the manufacturers’ protocol and were stored at -80°C.

#### In vitro binding

1-2 μg His-tagged calcineurin was first bound to magnetic Dynabeads (Thermo Fisher Sci. USA) in base buffer (50 mM Tris-HCl pH 7.5, 150 mM NaCl, 0.1% Tween 20, 1 mM β-mercaptoethanol, protease inhibitors, 5-10 mM imidazole, 1 mg/ml BSA) for 1 h at 4°C. 50-100 μg of bacterial cell lysate containing GST-peptide was then added to the binding reaction and incubated further for 2-3 hr. 3% of the reaction mix was removed as ‘input’ prior to the incubation, boiled in 2X-SDS sample buffer and stored at -20°C. The beads were washed in base buffer containing 15-20 mM imidazole. Bound proteins were then extracted with 2X-SDS sample buffer by boiling for 5 min. The proteins were analyzed by SDS-PAGE and immunoblotting with anti-GST (BioLegend MMS-112P) and anti-His (Qiagen 34660) antibodies. Blots were imaged with the Li-Cor Odyssey imaging system. GST peptides co-purifying with HIS-CN were normalized to their respective input and amount of calcineurin pulled down. Co-purification with CN was reported relative to that of the peptide with the known PxIxIT motif from NFATC1: PALES**PRIEIT**SCLGL. Statistical significance was determined with unpaired Student’s T test, using GraphPad. For Fig. S4B, FAM126A peptides used were FAM126A PSISIT: SGQQRP**PSISIT**LSTD and FAM126A ASASAA Mut: SGQQRP**ASASAA**LSTD.

### Hydrogen-Deuterium Exchange analysis (HDX-MS)

#### Protein expression

GST-tagged human calcineurin A (residues 2-391 of human CNA alpha isoform) in tandem with calcineurin B subunit were expressed in BL21 C41 *Escherichia coli*, induced with 0.1 mM IPTG (isopropyl β-d-1-thiogalactopyranoside) and grown at 23 °C overnight. Cells were harvested, flash frozen in liquid nitrogen, and stored at -80°C until use. Bacmids harboring MultiBac PI4KA complex constructs were transfected into *Spodoptera frugiperda* (Sf9) cells, and viral stocks amplified for one generation to acquire a P2 generation final viral stock. Final viral stocks were added to Sf9 cells at ∼ 1.8 × 10^6^ in a 1/100 to 1/50 virus volume to cell volume ratio. Constructs were expressed for 68 hrs before pelleting of infected cells. Cell pellets were snap frozen in liquid nitrogen, followed by storage at − 80 °C.

#### Protein purification (GST tagged human calcineurin)

*Escherichia coli* cell pellets were lysed by sonication for 5 min in lysis buffer [50 mM Tris pH 8.0, 100 mM NaCl, 2 mM EDTA, 2 mM EGTA, protease inhibitors (Millipore Protease Inhibitor Cocktail Set III, Animal-Free)]. NaCl solution was added to 1 M and the solution was centrifuged for 10 min at 12,000 × *g* at 1°C and for 20 min at 38,000 x *g* at 1°C (Beckman Coulter Avanti J-25I, JA 25.50 rotor). CHAPS was added to 0.02%. Supernatant was loaded onto a 5 ml GSTrap 4B column (GE) in a superloop for 45 min and the column was washed in Wash Buffer [50 mM Tris pH 8.0, 110 mM KOAc, 2 mM MgOAc, 1 mM DTT, 5% glycerol (v/v), 0.02% chaps] to remove nonspecifically bound proteins. The column was washed in Wash Buffer containing 2 mM ATP to remove the GroEL chaperone. The GST-tag was cleaved by adding Wash Buffer containing PreScission protease to the column and incubating overnight at 4 °C. Cleaved protein was eluted with Wash Buffer. Protein was concentrated using an Amicon 10 kDa MWCO concentrator (MilliporeSigma) and size exclusion chromatography (SEC) was performed using a Superdex 75 10/300 column equilibrated in Wash Buffer. Fractions containing protein of interest were pooled, concentrated, flash frozen and stored at − 80 °C.

#### Protein purification (PI4KA complex)

Sf9 pellets were resuspended in lysis buffer [20 mM imidazole pH 8.0, 100 mM NaCl, 5% glycerol, 2 mM βMe, protease (Protease Inhibitor Cocktail Set III, Sigma)] and lysed by sonication. Triton X-100 was added to 0.1% final, and lysate was centrifuged for 45 min at 20,000 x *g* at 1°C. (Beckman Coulter Avanti J-25I, JA 25.50 rotor). Supernatant was loaded onto a HisTrap FF Crude column (GE Healthcare) and superlooped for 1 h. The column was washed with Ni-NTA A buffer [20 mM imidazole pH 8.0, 100 mM NaCl, 5% glycerol (v/v), 2 mM βMe], washed with 6% Ni-NTA B buffer [30 mM imidazole pH 8.0, 100 mM NaCl, 5% (v/v) glycerol, 2 mM βMe], and the protein eluted with 100% Ni-NTA B buffer (450 mM imidazole). Elution fractions were passed through a 5 ml StrepTrapHP column pre-equilibrated in GF buffer [20 mM imidazole pH 7.0, 150 mM NaCl, 5% glycerol (v/v), 0.5 mM TCEP]. The column was washed with GF buffer before loading a tobacco etch virus protease containing a stabilizing lipoyl domain (Lip-TEV), and cleavage proceeded overnight. Cleaved protein was eluted with GF buffer and concentrated down to 250 µl in an Amicon 50 kDa MWCO concentrator (MilliporeSigma) pre-equilibrated in GF buffer. Concentrated protein was flash frozen in liquid nitrogen and stored at -80°C.

### Hydrogen-Deuterium Exchange Mass Spectrometry

#### Sample preparation

HDX reactions for PI4KA complex (PI4KIIIα, TTC7B, FAM126A) and Calcineurin were conducted in a final reaction volume of 24 µl with a final concentration of 0.17 µM (8 pmol) PI4KA complex and 0.95 µM (24 pmol) Calcineurin. The reaction was initiated by the addition of 16 µl of D_2_O buffer (150 mM NaCl, 20 mM pH 8.0 Imidazole, 90% D_2_O (V/V)) to 6.5 µl of PI4KA or PI4K buffer solution and 0.66 µl Calcineurin or Calcineurin buffer solution (final D_2_O concentration of 65%). The reaction proceeded for 3, 30, 300, or 3000 s at 20°C, before being quenched with ice cold acidic quench buffer, resulting in a final concentration of 0.6M guanidine-HCl and 0.9% formic acid post quench. All conditions and timepoints were generated in triplicate. Samples were flash frozen immediately after quenching and stored at -80°C until injected onto the ultra-performance liquid chromatography (UPLC) system for proteolytic cleavage, peptide separation, and injection onto a QTOF for mass analysis, described below.

#### Protein digestion and MS/MS data collection

Protein samples were rapidly thawed and injected onto an integrated fluidics system containing a HDx-3 PAL liquid handling robot and climate-controlled (2°C) chromatography system (LEAP Technologies), a Dionex Ultimate 3000 UHPLC system, as well as an Impact HD QTOF Mass spectrometer (Bruker). The full details of the automated LC system are described in ^75^. The protein was run over one immobilized pepsin column (Trajan; ProDx protease column, 2.1 mm x 30 mm PDX.PP01-F32) at 200 µL/min for 3 minutes at 10°C. The resulting peptides were collected and desalted on a C18 trap column (Acquity UPLC BEH C18 1.7mm column (2.1 x 5 mm); Waters 186003975). The trap was subsequently eluted in line with an ACQUITY 1.7 μm particle, 100 × 1 mm^2^ C18 UPLC column (Waters), using a gradient of 3-35% B (Buffer A 0.1% formic acid; Buffer B 100% acetonitrile) over 11 minutes immediately followed by a gradient of 35-80% over 5 minutes. Mass spectrometry experiments acquired over a mass range from 150 to 2200 m/z using an electrospray ionization source operated at a temperature of 200C and a spray voltage of 4.5 kV.

#### Peptide identification

Peptides were identified from the non-deuterated samples of PI4K using data-dependent acquisition following tandem MS/MS experiments (0.5 s precursor scan from 150-2000 m/z; twelve 0.25 s fragment scans from 150-2000 m/z). MS/MS datasets were analyzed using PEAKS7 (PEAKS), and peptide identification was carried out by using a false discovery-based approach, with a threshold set to 1% using a database of known contaminants found in Sf9 cells and BL21 C41 *Escherichia coli* ^76^. The search parameters were set with a precursor tolerance of 20 ppm, fragment mass error 0.02 Da, charge states from 1-8, leading to a selection criterion of peptides that had a -10logP score of 35.4 and 29.3 for the PI4KA complex and calcineurin, respectively.

#### Mass Analysis of Peptide Centroids and Measurement of Deuterium Incorporation

HD-Examiner Software (Sierra Analytics) was used to calculate the level of deuterium incorporation into each peptide. All peptides were manually inspected for correct charge state, correct retention time, and appropriate selection of isotopic distribution. Deuteration levels were calculated using the centroid of the experimental isotope clusters. Results are presented as relative levels of deuterium incorporation, with no correction for back exchange. The only correction was for the deuterium percentage of the buffer in the exchange reaction (65%). Differences in exchange in a peptide were considered significant if they met all three of the following criteria: ≥5% change in exchange, ≥0.5 Da difference in exchange, and a two-tailed T-test value of less than 0.01. The raw HDX data are shown in two different formats. The raw peptide deuterium incorporation graphs for a selection of peptides with significant differences are shown in Fig. 4e+g, with the raw data for all analyzed peptides in the source data (Supplementary Table 3). To allow for visualization of differences across all peptides, we utilized number of deuteron difference (#D) plots (Fig. 4d). These plots show the total difference in deuterium incorporation over the entire H/D exchange time course, with each point indicating a single peptide. Samples were only compared within a single experiment and were never compared to experiments completed at a different time with a different final D_2_O level. The data analysis statistics for all HDX-MS experiments are in Supplementary Table 2 according to published guidelines ^77^.The mass spectrometry proteomics data have been deposited to the ProteomeXchange Consortium via the PRIDE partner repository ^78^ with the dataset identifier PXD025900.

### *In vivo* analysis of FAM126A phosphorylation status

HeLa cells seeded onto 10 cm plates were transfected with EFR3B HA_T2A_ TTC7B MYC_P2A_FLAG FAM126A WT or ASASAA mutant. 24 h post-transfection, plates were washed, trypsinized and passaged onto 60 mm plates for treatments. The next day, cells were pre-treated with 2 μM FK506 (LC Laboratories) for 1 h, 2 μM BIM (bisindolylmaleimide, Calbiochem) for 15 min or DMSO in growth media. Cells were then stimulated with 500 nM PMA (Phorbol 12-myristate 13-acetate, Sigma-Aldrich) or DMSO for 15 min, washed and harvested in ice-cold PBS. Pellets were snap-frozen in liquid nitrogen and stored at -80°C until use. Cells were lysed with RIPA buffer (150 mM NaCl, 1% Triton X-100, 0.5% Deoxycholate, 0.1% SDS, 50 mM Tris pH 8.0) supplemented with protease and phosphatase inhibitor cocktail and 250U/ml benzonase for 30 min rotating end-over-end at 4°C and subjected to fine needle aspiration with sterile 27.5-gauge needle. Cell lysates were clarified by centrifugation (16,000g for 20min). Solubilized protein concentration in the supernatant were quantified using BCA analysis. 20 μg from each lysate was run out on 7.5% SDS-PAGE gels followed by analysis with Western blot. Changes in electrophoretic mobility of FAM126A were assessed by immunoblotting with anti-FLAG and custom anti-phosphospecific FAM126A S485 antibody. PKC activation was assessed by phosphorylation of the downstream substrate, ERK using p44/42 Erk1/2 antibody (1:3,000; 3A7, Cell Signaling Technologies). Phosphorylation status of FAM126A in each treatment was quantified as pFAM126A S485 signal/FLAG signal and reported relative to that of in DMSO treated FAM126A WT sample, using ImageStudio imaging software. Statistical significance was determined using GraphPad.

### Bioluminescence resonance energy transfer (BRET) measurements

HEK 293T cells were trypsinized and plated on white 96-well plates at a density of 75,000 cells/100 μl per well, together with the indicated DNA constructs (0.15 μg total DNA in 25 μl per well) and the cell transfection reagent (1.5 μl GeneCellin (Bulldog Bio) in 25 μl per well) in Opti-MEM reduced serum medium (Gibco). Cells were transfected with DNA encoding the human M3 muscarinic receptor (0.1 μg total DNA/well) and the previously established SidM-2XP4M-based PI4P biosensor ^28^ (0.05 μg total DNA/well). After 6 h, media were replaced with 100 μl/well Dulbecco/s modified Eaglès medium (DMEM) supplemented with 10% fetal bovine serum, 50 U/ml penicillin and 50 μg/ml streptomycin. Measurements were performed 28 h post-transfection. Prior to measurements, media of cells were replaced with 50 μl buffer containing 120 mM NaCl, 4.7 mM KCl, 1.2 mM CaCl_2_, 0.7 mM MgSO_4_, 10 mM glucose and 10 mM Na-HEPES, pH 7.4. Cells were pretreated with FK506 (1 μM), Cyclosporin A (10 μM) or DMSO for 1 h at 37°C. Measurements were performed at 37°C using a Varioskan Flash Reader (Thermo Scientific) and initiated with the addition of the cell permeable luciferase substrate, coelenterazine h (20 μl, final concentration of 5 μM). Counts were recorded using 485 and 530 nm emission filters. Detection time was 250 ms for each wavelength. The indicated reagents were also dissolved in modified Krebs-Ringer buffer and were added manually in 10 μl. For this, plates were unloaded, which resulted in an interruption in the recordings. All measurements were done at least in triplicates. BRET ratios were calculated by dividing the 530 nm and 485 nm intensities and results were normalized to the baseline. Since the absolute initial ratio values depended on the expression of the sensor, the resting levels were considered as 100%, whereas the 0% was determined from values of those experiments where cytoplasmic Renilla luciferase construct was expressed alone.

### Statistical Analysis

Statistics were computed using Graphpad Prism 9. All data shown as representative images or as the mean of measurements with standard deviation (SD) error bars unless noted otherwise. Data represent at least three independent experiments. For image analysis in Fig. 1f and Supplementary Fig.1f, number of cells analyzed for GM130 co-localization were as follows: n=166 for CNAβ2, n=164 for CNAβ1, n=130 for CNAβ1^C483^, n=128 for CNAβ1^C493S^, n=119 for CNAβ1^C2S^. For plasma membrane signal ratio measurements: n=75 for CNAβ2, n=86 for CNAβ1, n=98 for CNAβ1^C483^, n=80 for CNAβ1^C493S^, n=77 for CNAβ1^C2S^. Image analysis for Fig. 2g were performed on n=76 for vector control, n=94 for wildtype ABHD17A, n=89 for ABHD17A S190A mutant. Fig. 1g immunoblot is representative of n=5 for EFR3B-FLAG, n=5 for FLAG-CNAβ1, n=3 for each CNAβ1 mutant (C483S, C493S, C2S). Two-tailed unpaired Student’s *t-*test was applied for statistical analyses between two groups. One-way analysis of variance (ANOVA) with appropriate multiple comparisons (all indicated in figure legends) were performed when comparing more than two groups.

### Data Availability

All AP-MS data have been deposited to ProteomeXchange through partner MassIVE with the following identifiers: PXD026809 and MSV000087664, respectively. The dataset can be accessed at ftp://MSV000087664@massive.ucsd.edu (password: calcineurin). HDX-MS proteomics data have been deposited to the ProteomeXchange via the PRIDE partner repository with the dataset identifier PXD025900. The source data for Figs. 1c-d,f-g; 2b-f; 3d-e; 4b-c; 5a-b; 6a-b,d and Supplementary Figs 1b-c,f; 2a-b; 3b-c; 4b-c are provided as a Source Data file. All other primary data that support the findings of this study are available from the corresponding author upon reasonable request.

## Acknowledgements

We thank Callie Preast Wigington for critical reading of the manuscript, Rachel Bond for generating most of the CNAβ plasmids used in this study and for initiating experiments to identify CNAβ-interacting proteins, Pin-Joe Ko for helpful discussions about quantitative image analysis, Jamin Hein for assistance in developing the phospho-specific FAM126A antibody, the Skotheim lab for cell lines, the De Camilli lab for plasmids and all members of the Cyert lab for their support and critical feedback. M.S.C. and I.U.T. are supported by grants from the National Institute of Health R01GM129236 and R35GM136243. J.E.B. acknowledges funding support from a Discovery grant from the Natural Science and Engineering Research Council of Canada (NSERC-2020-04241, JEB), and the Michael Smith Foundation for Health Research (JEB, Scholar Award 17686). G.G. and T.B. are supported by the Intramural Research Program of the Eunice Kennedy Shriver National Institute of Child Health and Human Development of the NIH. A.C.G. is supported by a grant from the Canadian Institutes of Health Research (FDN 143301). P.V. is supported by the Hungarian National Research, Development and Innovation Fund (NKFIK134357). E.C. acknowledges support from the Canadian Institutes of Health Research (PJT-162184).

## Author contributions

I.U.T. and M.S.C. jointly designed the study. I.U.T. performed the majority of the experiments and analyzed data, supervised by M.S.C., with the exception of the following: M.A.H.P., M.L.J. and J.E.B. designed and executed HDX-MS experiments. J.R. carried out *in vitro* binding studies. A.Z.L.S. and E.C. designed and performed pulse-chase analyses of palmitate incorporation. N.S. and A.C.G. designed and conducted AP-MS experiments. BRET-based analyses of PI4P dynamics were designed and executed by P.V. (effects of CN inhibitors) and by G.G. and T.B. (analyses of FAM126A mutants). I.U.T. and M.S.C. wrote the manuscript with editorial input and approval from all authors. Correspondence and requests for materials/plasmids should be addressed to M.S.C.

