## Supplementary Information for "Palmitoylation targets the Calcineurin phosphatase to the Phosphatidylinositol 4-kinase complex at the plasma membrane"

**Legends for Supplemental Figures 1- 5:**

**Supplementary Figure 1:** (related to Fig. 1) **a** Representative IF images of HeLa cells expressing FLAG-CNAβ1 WT or CNAβ1 C483/C493S double mutant. Scale bar = 15 μm **b** Detection of palmitoylation using Acyl-RAC in COS-7 cells transfected with FLAG-CNAβ2, FLAG-CNAβ1 (WT or cysteine mutants) or EFR3B-FLAG. **c** Analysis of CNAβ1 palmitoylation in the cytosol and membrane fractions of COS-7 cells labelled with 17-ODYA. Immunoblot illustrates total CNAβ1 using anti-GFP and 17-ODYA detected using streptavidin following CLICK chemistry with azide-Biotin. Level of CNAβ1 palmitoylation quantified by the streptavidin signal (17-ODYA) / total protein signal (GFP). Data are mean ± SD (n=2). **d** Representative image of the 5-pixel wide region of interest (ROI) generated at the cell periphery to define the PM of each image quantified in Fig.1e and Supplementary Fig. 1f. **e** Representative IF images of COS-7 cells expressing FLAG-CNAβ1 C483S (top panel) or C493S (bottom panel) with PM-marker Venus-RIT (green) immunostained with anti-FLAG (red) and anti-GM130 antibodies (blue). Top panel of CNAβ1 C483S is an example of a cell with weak localization to the PM (green box) and Golgi (red box). Scale bar = 15 μm **f** Co-localization of FLAG-CNAβ mutants with GM130 quantified using EZ colocalization plug-in in Image J. Data are presented as median Pearson`s coefficients. At least 100 cells were analyzed across three replicates. *p <0.05, **** *p* <0.0001 using one-way ANOVA followed by Kruskal-Wallis test. Bottom graph: anti-FLAG signal intensity at the cell periphery (as in d) normalized to total intensity. Data are presented as median. At least 70 cells were imaged across three replicates. * *p* <0.05, **** *p* <0.0001 using one- way ANOVA followed by Kruskal-Wallis test.

**Supplementary Figure 2:** (related to Fig. 2) **a** GFP-CNAβ1 palmitoylation levels in COS-7 cells co-expressing vector or mcherry-APT1 determined using metabolic labelling with 17-ODYA. **b** GFP-CNAβ1 palmitoylation levels in **a** quantified by the streptavidin signal (17-

ODYA) / total protein signal (GFP) normalized to vector control. Data are mean ± SEM (n=3). n.s. not significant.

**Supplementary Figure 3:** (related to Fig. 3) **a** Complete dotplot of CN interactor proteins identified by AP-MS analysis. **b** Immunoblot analysis of the anti-GFP immunoprecipitates from HEK 293 inducible cell lines expressing GFP vector, GFP-CNA$\alpha$, GFP-CNAβ2 or GFP-CNAβ1, transfected with FLAG-NFAT2. **e** Amount of FLAG-NFAT2 co-purified with GFP across samples quantified as bound FLAG signal/ bound GFP signal normalized to input. CNB interaction was quantified as bound anti-CNB signal/ bound GFP signal normalized to input. Data are mean of ± SEM (n=3), relative to GFP-CNA$\alpha$. n.s. not significant, ** $p < 0.01$ using one-way ANOVA with Dunnette's multiple comparison tests. **d** Schematic diagram of the expression plasmid harboring DNA sequences encoding EFR3B-HA (red), TTC7B-MYC (or GFP, green) and FLAG-FAM126A (blue) separated by 2A viral peptides, T2A and P2A (gray) used in this study. A separate plasmid encoding HA or GFP tagged PI4KA was used when necessary. **e** Representative images of fixed HeLa cells transfected with the plasmid in **d** immunostained with anti-HA, anti-MYC or anti-FLAG. Scale bar = 15 μm. **f** Immunoblot of HeLa lysate expressing the 2A plasmid shown in **(d),** arrow shows residual uncut P2A form.

**Supplementary Figure 4:** (related to Fig. 4) **a** Sequence alignment depicting the evolutionary conservation of the FAM126A C-terminus (a.a 480-521) across vertebrates. CN binding PxIxIT motif: PSISIT (red) and phospho-serine 485 identified in this study (yellow) are boxed. **b** Representative immunoblot showing co-purification of GST-tagged peptides from FAM126A containing WT or mutated PxIxIT motif (WT: SGQQRPPSISITLSTD or ASASAA mut: SGQQRPASASAALSTD) with His-CN heterodimer (CN WT or CN NIR). **c** Quantification of anti-HIS co-purifications as in **b** relative to that of the peptide with the PxIxIT motif from NFATC1. Data are mean ± SEM; (n=3). n.s. not significant, ** $p < 0.01$ **d**

Deuterium incorporation difference for all peptides that showed a significant increase or decrease in exchange (>5%, 0.4 Da, and an unpaired t-test $p$ <0.01). For all panels, error bars show SD (n = 3). **e** The number of HDX for all analyzed peptides over the entire time course. Every point represents the central residue of an individual peptide. Significant peptides are highlighted in red. Error bars show SD (n=3).

**Supplementary Figure 5: a** Immunoblot showing electro-mobility shifts in FAM126A$^{ASASAA}$ in lysates of HeLa cells co-expressing HA-PI4KA and TTC7B-GFP in the presence or absence of EFR3B-HA. PI and PII denote phosphorylated forms, deP denotes dephosphorylated FAM126A. **b** Representative images of fixed HeLa cells transfected with EFR3B HA_T2A_TTC7 MYC_P2A_ FLAG FAM126A (WT or ASASAA) plasmids**,** stained with anti-FLAG (red), anti-pFAM126A S485 (green) antibodies and DAPI (blue). Scale bar = 15 µm. **c** Images showing localizations of GFP-PI4K (green) and FLAG-FAM126A (WT or ASASAA mutant) in COS-7 cells co-expressing EFR3B-HA, TTC7B-MYC. Representative images of cells fixed and immunostained with anti-FLAG (red) and anti-GM130 (blue) antibodies are shown. Scale bar = 15 µm

**Additional information is presented in supplemental tables (1-3)**

**Supplementary Table 1:** *High confidence interactors identified by AP-MS results.* See attached tabular data file.

**Supplementary Table 2:** *HDX-MS data analysis statistics summary.*

| Protein Data Set | PI4KA | FAM126A | TTC7B | Calcineurin A | Calcineurin B |
|---|---|---|---|---|---|
| HDX reaction details | %D2O=65% pH(read)= 7.5 Temp= 18°C | %D2O=65% pH(read)= 7.5 Temp= 18°C | %D2O=65% pH(read)= 7.5 Temp= 18°C | %D2O=65% pH(read)= 7.5 Temp= 18°C | %D2O=65% pH(read)= 7.5 Temp= 18°C |
| HDX time course | 3s, 30s, 300s, 3000s | 3s, 30s, 300s, 3000s | 3s, 30s, 300s, 3000s | 3s, 30s, 300s, 3000s | 3s, 30s, 300s, 3000s |
| HDX controls | N/A | N/A | N/A | N/A | N/A |
| Back-exchange | Corrected based on %D2O | Corrected based on %D2O | Corrected based on %D2O | Corrected based on %D2O | Corrected based on %D2O |
| Number of peptides | 234 | 53 | 106 | 81 | 40 |
| Sequence coverage | 77.6% | 80.9% | 84.2% | 89% | 89.3% |
| Average peptide length / Redundancy | Length = 13.3 Redundancy = 1.3 | Length = 14.4 Redundancy = 1.3 | Length = 12.3 Redundancy = 1.3 | Length = 11.5 Redundancy = 2.3 | Length = 14.0 Redundancy = 3.2 |
| Replicates | 3 (3s apo and 30s complex in duplicate) | 3 (3s apo and 30s complex in duplicate) | 3 (3s apo and 30s complex in duplicate) | 3 (30s complex in duplicate) | 3 (30s complex in duplicate) |
| Repeatability | Average StDev = 0.7% | Average StDev = 0.8% | Average StDev = 0.6% | Average StDev = 0.8% | Average StDev = 0.6% |
| Significant differences in HDX | >5% and >0.5 Da and unpaired t-test <0.01 | >5% and >0.5 Da and unpaired t-test <0.01 | >5% and >0.5 Da and unpaired t-test <0.01 | >5% and >0.5 Da and unpaired t-test <0.01 | >5% and >0.5 Da and unpaired t-test <0.01 |

**Supplementary Table 3:** *HDX-MS source data.* Raw data for all analyzed peptides with protein coverage maps for PI4KA, TTC7B, FAM126A, CNA/CNB. See attached tabular data file.

Supplementary Figure 1

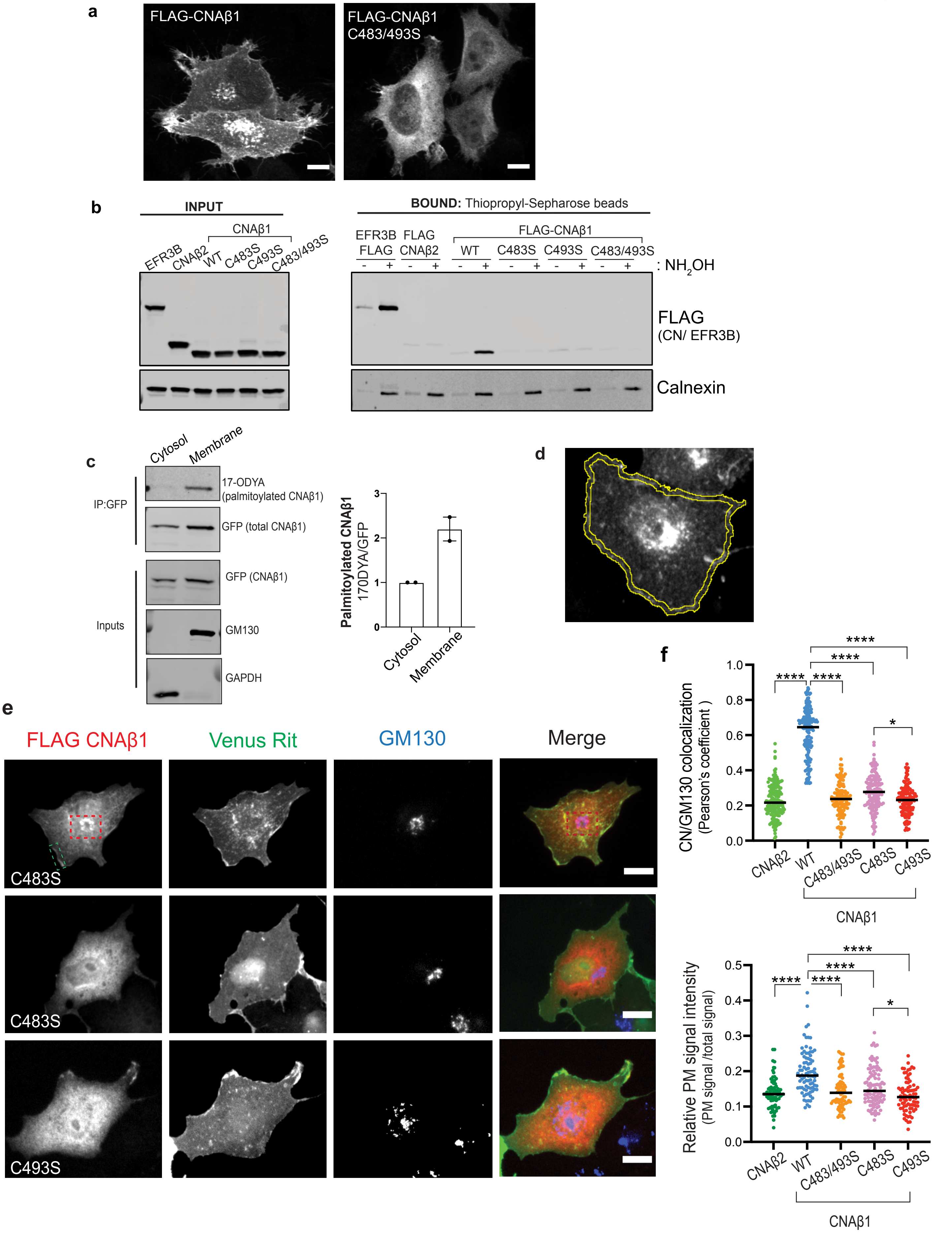

Supplementary Figure 2

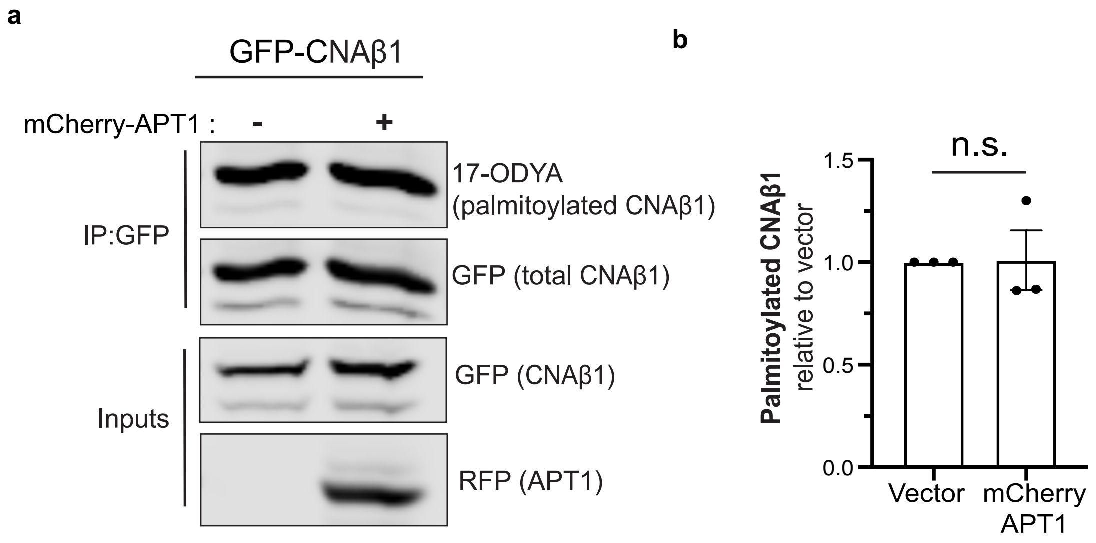

Supplementary Figure 3

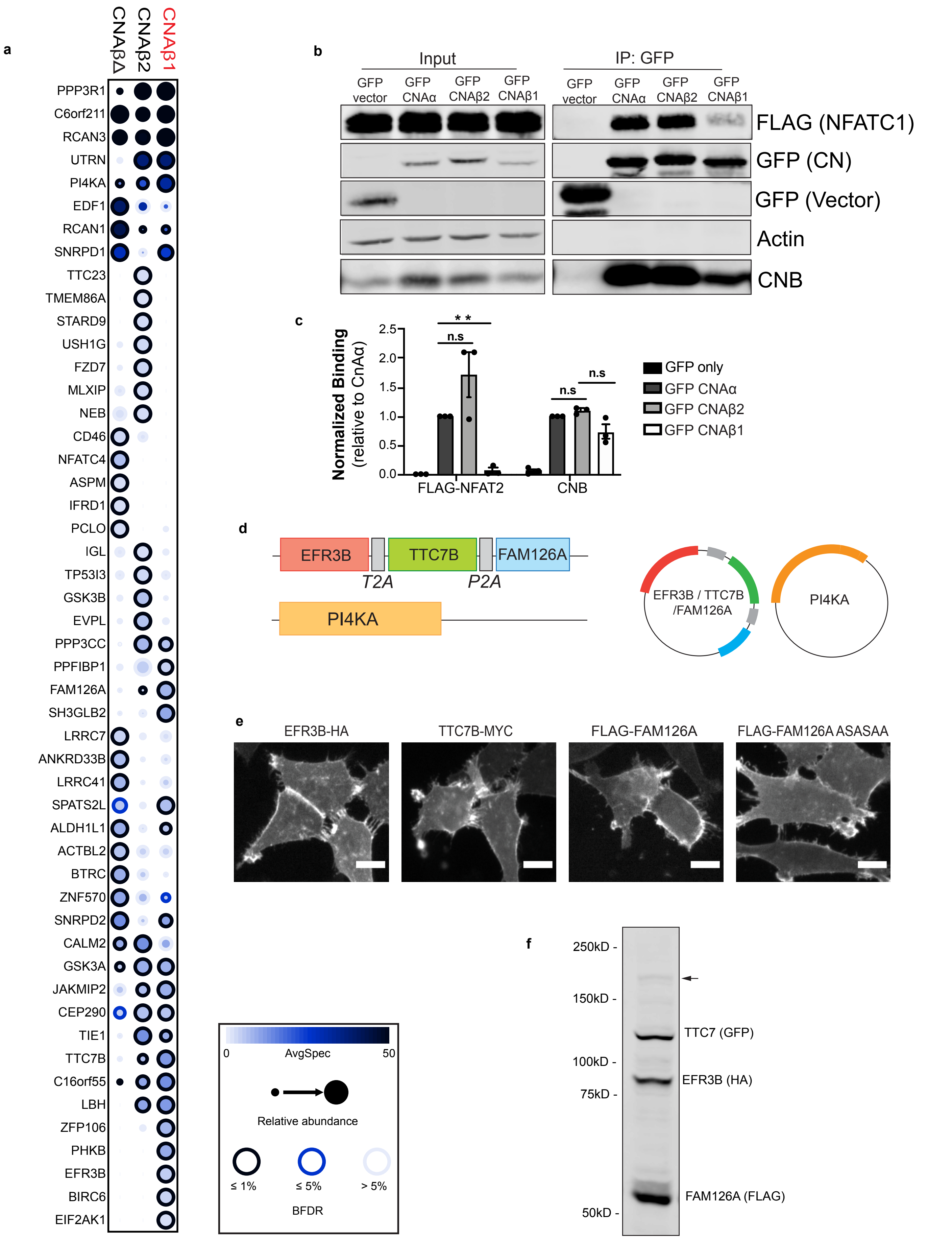

### Supplementary Figure 4

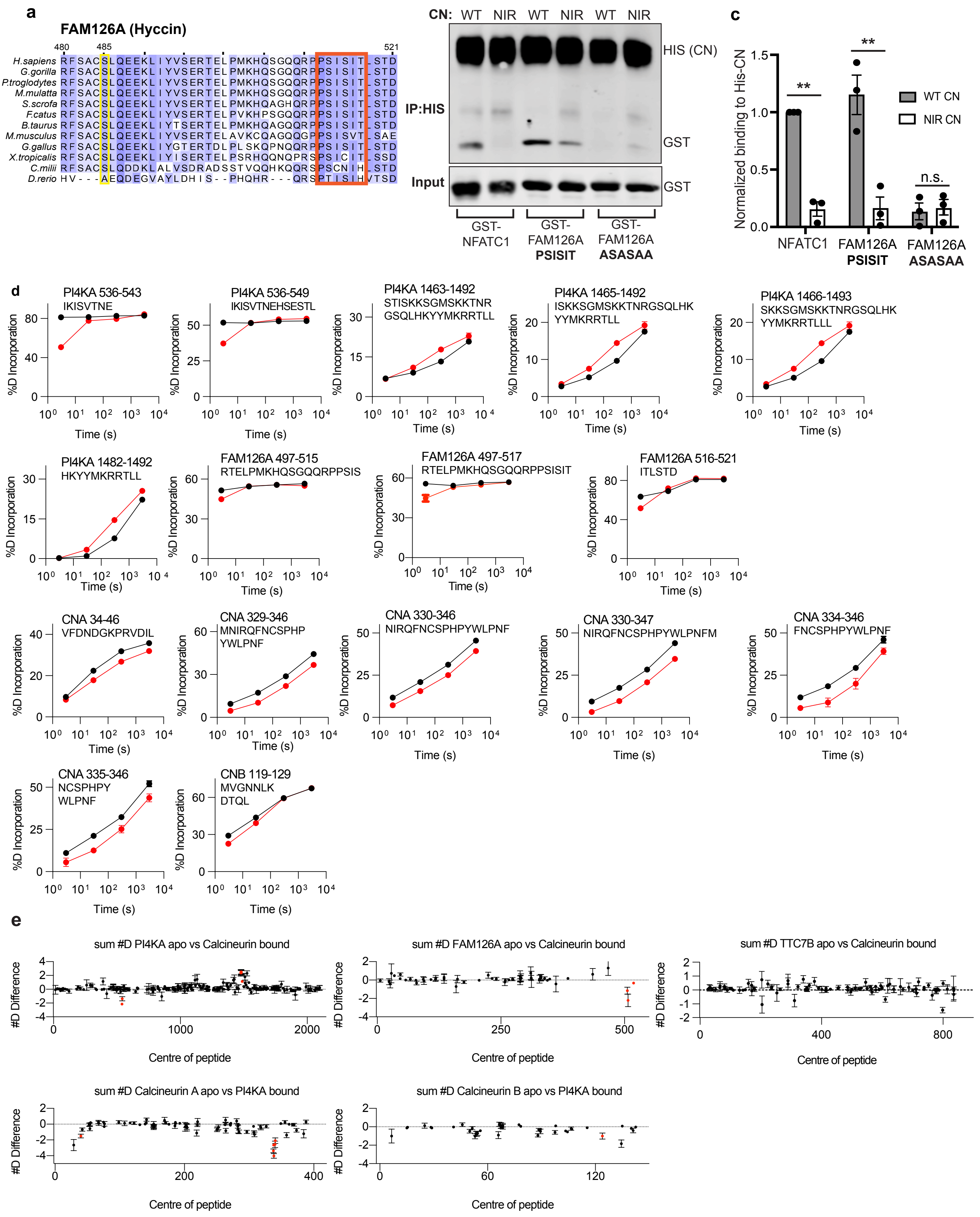

### Supplementary Figure 5

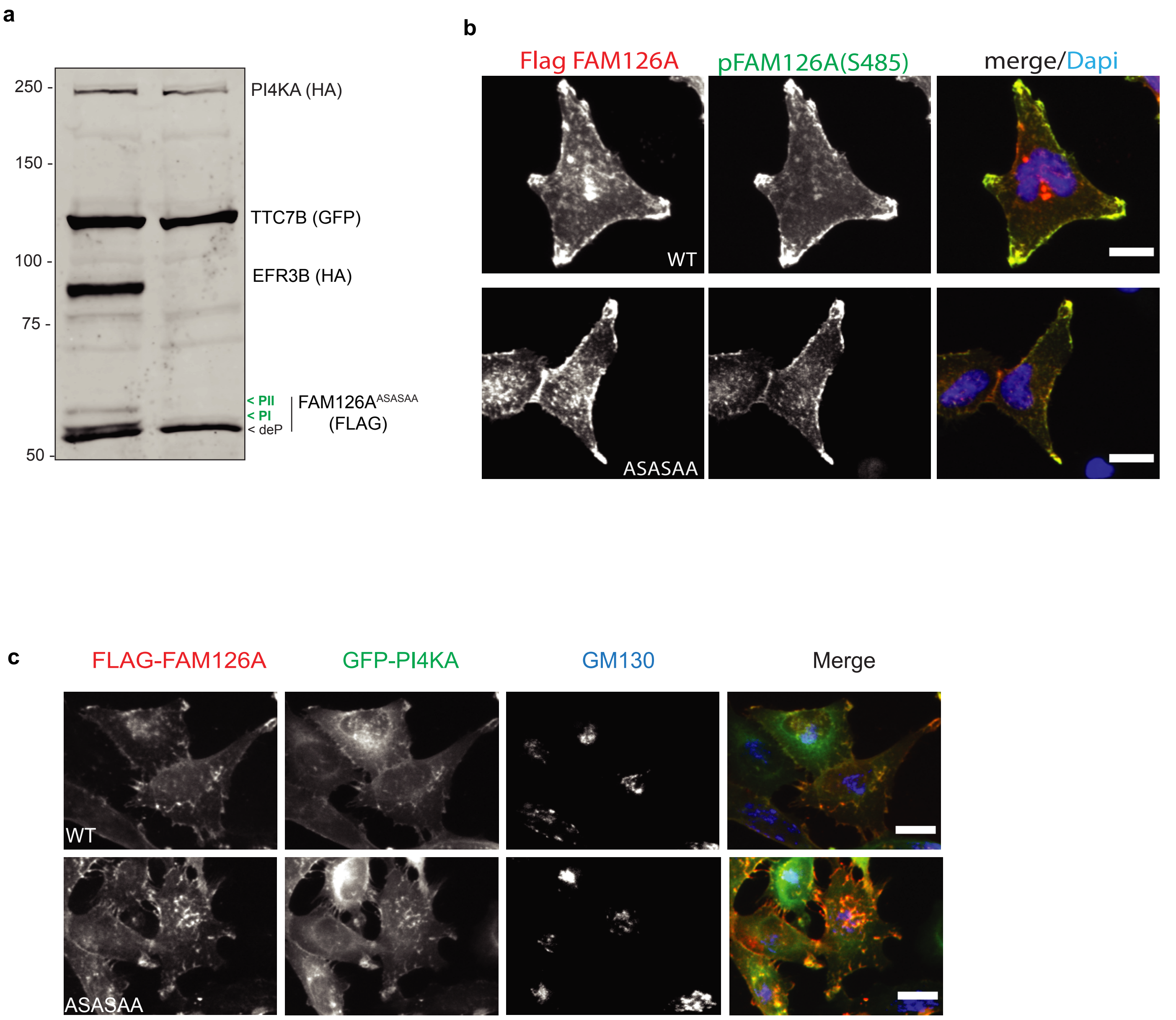
